# Allosteric HSP70 inhibitors perturb mitochondrial proteostasis and overcome proteasome inhibitor resistance in multiple myeloma

**DOI:** 10.1101/2020.04.21.052456

**Authors:** Ian D. Ferguson, Yu-Hsiu T. Lin, Christine Lam, Hao Shao, Martina Hale, Kevin M. Tharp, Margarette C. Mariano, Veronica Steri, Donghui Wang, Paul Phojanokong, Sami T. Tuomivaara, Byron Hann, Christoph Driessen, Brian Van Ness, Jason E. Gestwicki, Arun P. Wiita

## Abstract

Proteasome inhibitor (PI) resistance remains a central challenge in multiple myeloma. To identify pathways mediating resistance, we first map proteasome-associated genetic co-dependencies. We identify cytosolic heat shock protein 70 (HSP70) chaperones as potential targets, consistent with proposed mechanisms of myeloma tumor cells overcoming PI-induced stress. These results lead us to explore allosteric HSP70 inhibitors (JG compounds) as myeloma therapeutics. We show these compounds exhibit increased efficacy against acquired and intrinsic PI-resistant myeloma models, unlike HSP90 inhibition. Surprisingly, shotgun and pulsed-SILAC proteomics reveal that JGs overcome PI resistance not via the expected mechanism of inhibiting cytosolic HSP70s, but instead through mitochondrial-localized HSP70, HSPA9, destabilizing the 55S mitoribosome. Analysis of myeloma patient data further supports strong effects of global proteostasis capacity, and particularly *HSPA9* expression, on PI response. Our results characterize dynamics of myeloma proteostasis networks under therapeutic pressure while motivating further investigation of HSPA9 as a specific vulnerability in PI-resistant disease.

## Introduction

Nearly all multiple myeloma (MM) patients receive a proteasome inhibitor (PI) as part of their therapeutic regimen^1^. In malignant plasma cells, treatment with PIs induces unfolded protein stress^2,3^ and it is thought that these cells are preferentially sensitive because of their extremely high levels of immunoglobulin synthesis. Specifically, misfolded immunoglobulins cannot be efficiently degraded when proteasomes are inhibited, leading to their accumulation within the endoplasmic reticulum (ER) and activation of apoptosis via the unfolded protein response (UPR)^4^. However, while the large majority of MM patients will initially respond to PI therapy, none will be cured. This inevitable relapse makes overcoming PI resistance a leading, and long-standing, conundrum for MM clinicians.

Currently, there are many proposed modes of resistance to PI therapy, including changes in the immune microenvironment and/or mutations in *PSMB5*, the proteasome subunit bound by PIs^5–7^. However, the leading model of PI resistance is that MM cells rewire protein homeostasis (proteostasis) to decrease unfolded protein stress. For example, MM cell lines and primary patient samples can become resistant by decreasing immunoglobulin synthesis^8^. Conversely, MM plasma cells can increase their ability to degrade or fold proteins, allowing them to adapt to PI inhibition. Indeed, analysis of *in vitro*-evolved, PI-resistant MM cells have shown upregulation of both proteasome subunits, including PSMB5, and protein-folding chaperones, most notably HSP70 and HSP27 isoforms^9^. In addition, some studies have proposed that downregulation of 19S proteasomal regulatory “cap” subunits may relate to PI resistance, possibly by altering which proteins are degraded^10,11^. Together, these observations suggest that one important mechanism of PI resistance is the compensatory tuning of proteostasis.

There are hundreds of proteins associated with proteostasis. Which of these might be targeted to overcome PI resistance? Our prior work^12^, as well as that of others^13^, found that acute PI treatment induces a heat shock response. This response leads to marked upregulation of the inducible cytosolic HSP70 (*HSPA1A* gene), as well as BAG co-chaperones, which are known to be required for HSP70’s activities. These observations suggest that targeting the HSP70 axis may partially eliminate a mechanism for acquiring PI resistance. In addition, PI-resistant cells might become selectively vulnerable to inhibitors of this chaperone.

While several small molecule HSP70 inhibitors have been developed, we focused on a class of “JG” compounds, including JG98, that allosterically inhibit HSP70 by disrupting its interaction with the BAGs^14–16^. Thus, these molecules target the same sub-network that was previously identified as being involved in resistance to PIs. Here, we investigate the potential of these “JG” series compounds to synergize with PI and overcome PI resistance in MM. The results of *in vitro* screening support the therapeutic potential of these agents to specifically target PI-resistant disease. This relationship seemed to be of particular importance, as inhibitors of other proteostasis proteins were not effective in this setting. To understand the mechanism of this interaction, we performed shotgun and pulsed-SILAC proteomics, uncovering an unexpected role for mitochondrial proteostasis. Overall, these results suggest a new approach to overcome PI-resistance in MM, while also revealing broader interactions between sub-cellular proteostasis networks in this paradigmatic disease.

## Results

### Cytosolic HSP70 shows strongest genetic co-dependency with proteasome subunits in genome-wide CRISPR screen data

To reveal potential mechanisms of PI resistance, we first interrogated genome-wide CRISPR-knockout screening data in the Cancer Dependency Map (DepMap)^17^. We specifically asked: if cancer cell lines are highly dependent on the proteasome for survival, what other proteostasis-related genes are they also dependent on? We reasoned that this “co-dependency” approach could allow us to identify genes that could either be favorable targets for PI combinations or to overcome PI resistance. For this analysis, we first manually curated a list of 441 proteostasis genes based on prior literature (**Supplementary Dataset 1**). We then evaluated the pairwise Pearson correlation of the survival dependency score for the 406 of these genes included in CRISPR screens across 558 cell lines in DepMap (Release 19Q1). We developed an overall landscape illustrating both positive (cells are sensitive to genetic depletion of both genes) and negative (cells sensitive to depletion of one gene tend not to be sensitive to depletion of the other) correlations (**Fig. 1A-B, Supplementary Fig. 1A)**

**Figure 1.**
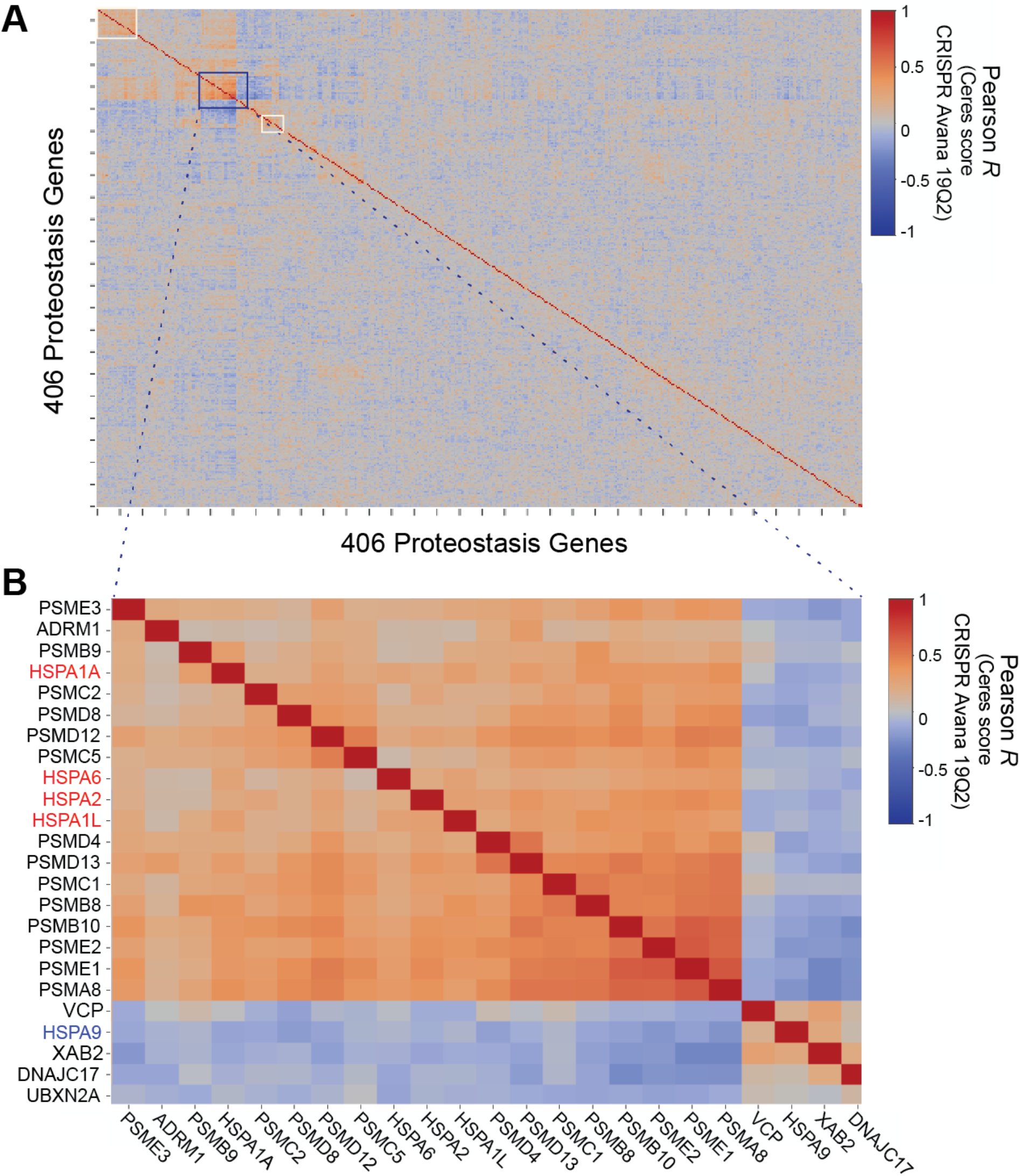
Cytosolic HSP70 shows strongest genetic co-dependency with proteasome subunits in genome-wide CRISPR screen data. **A.** 406 genes involved in protein homeostasis were used in a Pearson Correlation clustering analysis of genome-wide pan-cancer CRISPR knockout dependency screen dataset downloaded from the DepMap portal (19Q1 release). **B.** The most prominent co-dependency cluster includes cytosolic HSP70s (red) and proteasome subunits. Additional clusters highlighted in white squares in **A.** analyzed in **Supplementary Fig. 1**.

We found that subunits of the proteasome formed the most notable clusters of positively correlated genes (**Fig. 1B**), as expected. However, the next most prominent genes were those encoding several cytosolic HSP70 homologs: *HSPA1A* (HSP72 protein), *HSPA6* (HSP70B), *HSPA2* (HSP70-2), *HSPA1L* (HSP70-1L). This result supports previous analyses in MM^12, 18^ underscoring that pharmacologic blockade of both the proteasome and cytosolic HSP70s might be synergistic. We also observed a smaller cluster of proteasome subunits which included the ER-localized HSP70, *HSPA5* (BiP/GRP78) (**Supplementary Fig. 1B**), suggesting this homolog as another promising target. Notably, the mitochondria-localized HSP70, *HSPA9* (MtHSP70/GRP75/mortalin), was anti-correlated with the proteasome clusters.

Another interesting result of this DepMap analysis was that very few other proteostasis pathways were linked to the proteasome. We did observe a co-dependency network involving several members of the ER-resident protein disulfide isomerase family (*P4HB*, *P4HA1*, *P4HA2*, *P4HA3*), a BiP-interacting co-chaperone (*DNAJC1*), and several other genes of diverse function (**Supplementary Fig. 1C**). While investigating these additional associations is beyond the scope of our work here, we have developed an interactive web-based tool (https://tony-lin.shinyapps.io/proteostasis-map/) for use by others in the proteostasis field. Together, these results suggest that the proteasome interaction with cytosolic, and potentially ER-localized HSP70, might be uniquely important.

### PI-resistant MM models show increased sensitivity to allosteric HSP70 inhibitors

Based on prior data and our analysis, we investigated the “JG” series of allosteric HSP70 inhibitors in MM models. The parent compound in this series, JG98, has been explored in other cancer models^19–21^. Recently, a medicinal chemistry campaign was used to improve the potency, stability and safety of this molecule^16^. To obtain an overview of the structure-activity relationships (SAR) in MM models, we chose a representative set of 16 analogs from this series (**Supplementary Fig. 2**). We evaluated the sensitivity of a well-characterized AMO-1 MM cell line evolved to be PI-resistant (AMO1-BtzR) and compared it to the parental counterpart (AMO1-WT)^9,22^ (**Fig. 2A**). We were very encouraged to find that 15 of 16 compounds demonstrated lower LC_50_’s in AMO1-BtzR vs. WT cells (**Fig. 2B-C**). We found this result particularly noteworthy as essentially all prior molecules described to overcome PI resistance in MM show approximately equal sensitivity between WT and PI-resistant cells (for example refs.^23–26^). The greater sensitivity of AMO1-BtzR cells to HSP70 inhibition supports a special dependency of PI-resistant cells on HSP70s.

**Figure 2.**
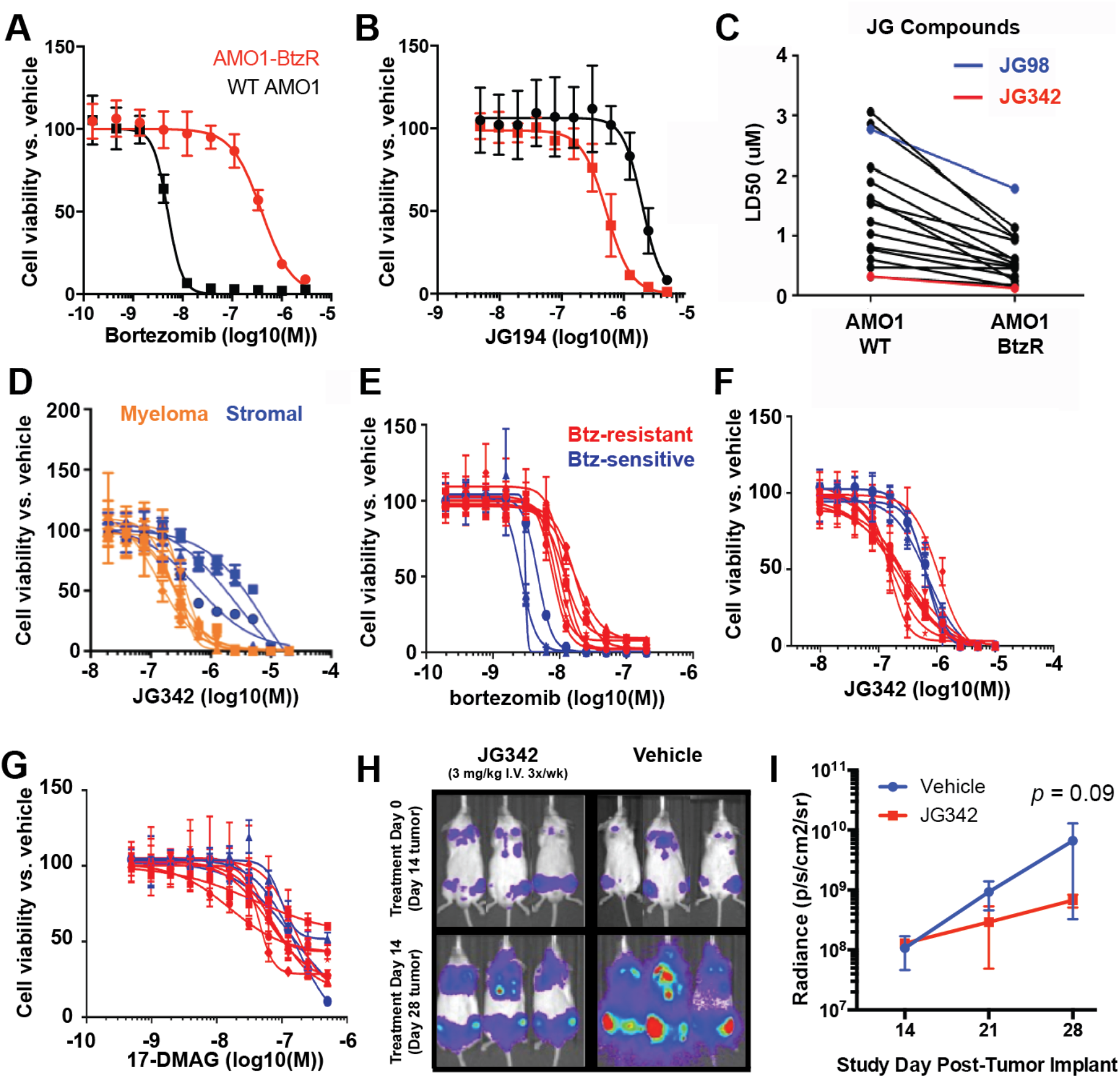
PI-resistant MM models show increased sensitivity to allosteric HSP70 inhibitors. **A-B.** AMO-1 bortezomib resistant (BtzR) cells are more sensitive than WT cells to an example JG compound, JG194. **C.** A larger panel of JG compounds (*n* =16) also show increased potency against AMO-1 BtzR MM model than WT. **D.** JG342 exhibits LC_50_’s in the nM range against a panel of MM cell lines and exhibits a therapeutic index against both immortalized (HS5, HS27A) and patient-derived bone marrow stromal lines. **E-G.** JG342 exhibits increased potency against MM cell lines with intrinsic resistance to Bortezomib. HSP90 inhibitor 17-DMAG does not show the same phenotype. All results in A-G. performed in quadruplicate in 384 well plates, with viability measured using CellTiterGlo at 48 hours. **H-I.** NSG mice (*n* = 3 per arm) were implanted with luciferase labeled RPMI-8226 MM cell line and dosed for two weeks with 3 mg/kg JG342 three times per week starting at day 14. JG342 exhibits modest in-vivo anti-MM activity, quantified in (**I.**). All error bars indicate +/− S.D.

Of the JG compounds tested, JG342 showed the lowest LC_50_ (122.6 nM) vs. AMO1-BtzR cells (**Fig. 2C**), leading us to focus on this molecule as a chemical probe for further mechanistic studies. First, we screened JG342 for activity against a panel of MM cell lines (MM.1S, RPMI-8226, U266, L363, AMO-1, KMS11, KMS34, JJN3) and compared these values to JG342 activity in non-malignant bone marrow stromal cells (immortalized HS5 and HS27A lines and one low-passage, patient-derived line) (**Fig. 2D**). We found that the MM cell lines were routinely more sensitive to JG342, suggesting a good therapeutic index.

We also expanded our analysis to MM cell lines recently defined in a large-scale screen to be intrinsically resistant (LP-1, KMS12-BM, MMM1, JIM-3) or sensitive (KMM1, MM-1144, KMS-18) to several different PIs, along with three evolved resistant lines (ANBL6-BtzR, RPMI-8226-BtzR, and U266-BtzR)^27^. We first confirmed the findings of this prior study, demonstrating that lines previously defined as PI resistant or sensitive showed the same phenotype in our hands (**Fig. 2E**). We next tested four of the JG compounds versus this cell line panel. Indeed, we found that the PI-resistant lines again showed increased sensitivity to JGs compared to their sensitive counterparts (**Fig. 2F, Supplementary Fig. 3A-C**).

We wondered whether the expression levels of HSP70s or BAGs might correlate with sensitivity across these MM and stromal cell lines. However, neither total HSP70 nor BAG3 expression was predictive of sensitivity to JG compounds (**Supplementary Fig. 3D**). We also wondered if inhibitors of other chaperones would also provide the same selective toxicity against PI-resistant cells or if this property was special to HSP70 inhibitors. We found that the HSP90 inhibitor, 17-DMAG, did not exhibit differential efficacy vs. PI-sensitive or resistant cells (**Fig. 2G**). Thus, the relationship between HSP70 and proteasome inhibition was not a general property of other proteostasis targets, consistent with the DepMap analysis.

Next, we evaluated the activity of JG342 in MM animal models *in vivo*. Prior to these experiments, we first determined the pharmacokinetics (PK) of JG342 in mice. After a single *i.v.* injection (3 mg/kg), the plasma concentrations decreased below the predicted therapeutic range (<150 nM) after ~6 hr (**Supplementary Fig. 3E**). This rate of clearance might be sufficient to give an anti-tumor effect if cell death is rapidly induced. To test this idea, we implanted luciferase-labeled RPMI-8226 cells intravenously into NOD *scid* gamma (NSG) mice, leading to disseminated disease in hematopoietic organs. After 14 days of tumor growth, we treated mice 3x/week for 2 weeks at 3 mg/kg IV JG342; by bioluminescent imaging we indeed noted a trend toward decreased tumor burden in treated mice vs. vehicle control (*p* = 0.09) (**Fig. 2H-I**).

However, we found that this treatment schedule led to a decrease in body weight (**Supplementary Fig. 3F**) and higher doses (5 mg/kg) led to rapid weight loss and mortality in a pilot cohort (data not shown). Thus, while our results suggest that targeting HSP70 carries significant promise to overcome PI resistance in MM, additional work is required to improve PK and safety.

### JG98 combination with proteostasis inhibitors leads to differential synergy and antagonism in MM cells

Although JG98 and its analogs were not immediately suitable for continued pre-clinical evaluation, we considered them useful chemical probes for further mechanistic studies. Accordingly, we sought to further investigate why PI-resistant MM cells are more sensitive to JG analogs. We also wanted to test whether targeting HSP70 in combination with PIs drives increased MM cell death.

Toward this latter point, previous work in MM models has suggested that simultaneously targeting other parts of the proteostasis network can be synergistic with PIs^25^. Extending from these findings, we aimed to measure the effects of inhibiting HSP70 at the same time as inhibiting three other, key proteostasis nodes: the proteasome, HSP90, and p97. In these experiments, we selected the representative inhibitors: bortezomib (PI), 17-DMAG (Hsp90 inhibitor)^28^, and CB-5083 (p97 inhibitor)^29^. We performed these combination treatment studies in three MM cell lines (RPMI-8226, MM.1S, KMS-34), as well as one acute myeloid leukemia (AML) cell line (CMK) to evaluate possible MM-specific effects. To inhibit HSP70 in these combinatorial studies we used JG98, the most well-characterized of the JG series. Cell proliferation was measured using CellTiterGlo and compound treatments were performed at the same time. Then, we assessed the effects of each pairwise drug combination on proliferation using the ZIP synergy score method^30^ (examples of raw viability data shown in **Fig. 3A-B**). By this metric, a score of zero is consistent with additivity, a positive score denotes synergy, a negative score antagonism, and the absolute value of the score reflects the strength of the interaction (**Fig. 3C**).

**Figure 3.**
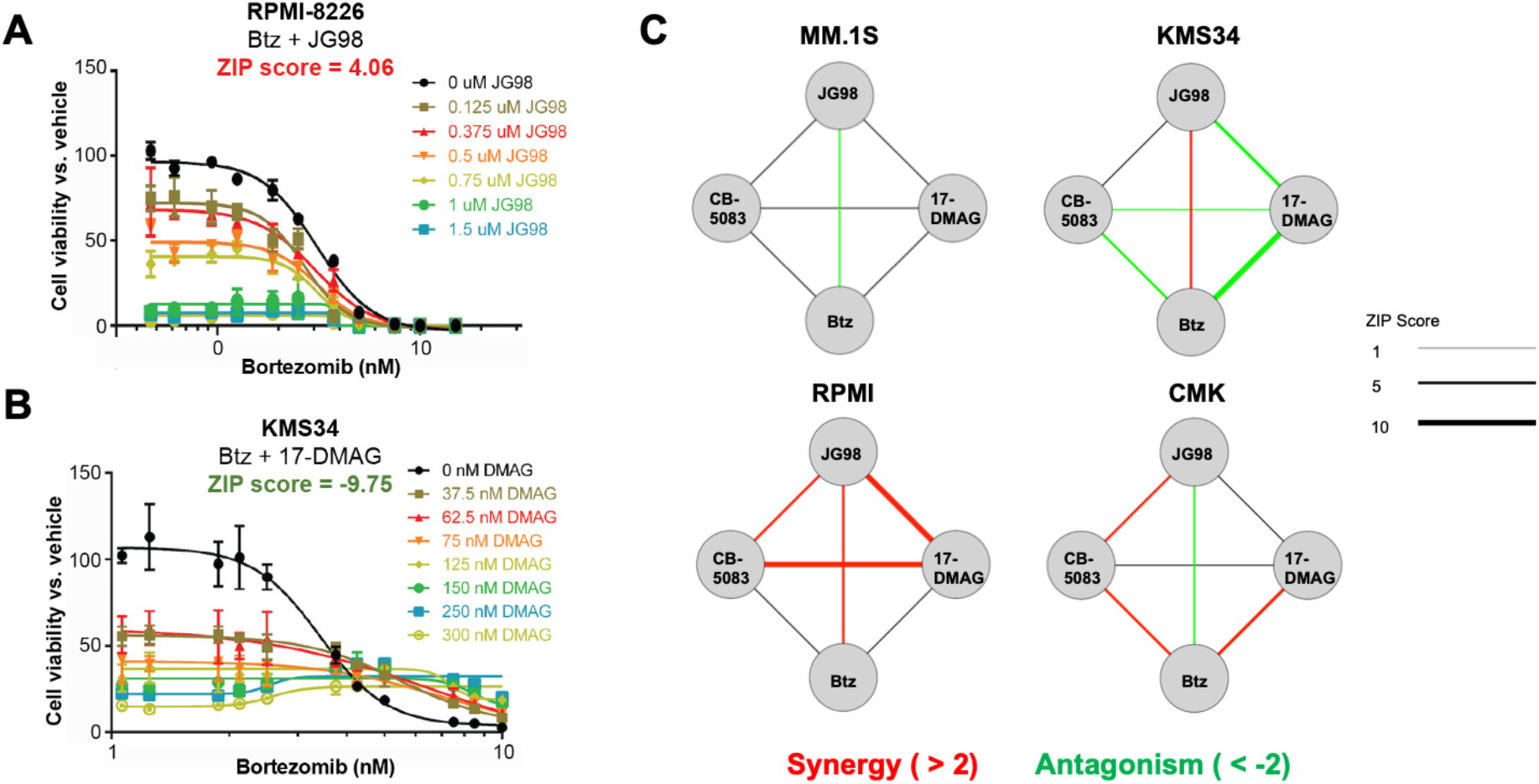
JG98 combination with proteostasis inhibitors leads to differential synergy and antagonism in MM cells. **A.** Bortezomib and JG98 combination shows mild synergy in RPMI-8226 myeloma cells as assessed by ZIP score. **B.** Bortezomib and 17-DMAG (HSP90 inhibitor) show antagonism in KMS34 cells. **C.** Network maps showing synergy and antagonism scores for pairwise drug screens as in (A-B) across three myeloma lines (MM.1S, KMS-34, and RPMI-8226) and one AML cell line (CMK). Thickness of the line denotes strength (as absolute value) of synergy or antagonism, and scores between −2 and 2 were considered additive.

Contrary to our hypothesis, we found that the JG98 plus bortezomib combination was essentially additive, with either weak synergy or antagonism noted across all four cell lines. Of the combinations tested, only the CB-5083 plus JG98 combination did not show any antagonism across all tested cell lines. For other combinations, we surprisingly found that synergy and antagonism was largely cell line specific (**Fig. 3C**, **Supplementary Fig. 4A**). For example, we noted strong synergy between JG98 and 17-DMAG in RPMI-8226 cells, but moderate antagonism in KMS-34. As one potential explanation for this finding, we examined available RNA-seq data from the Cancer Cell Line Encyclopedia (CCLE) to assess whether transcript expression of each target or their interaction partners may offer some explanation for these differential combination effects. However, we found no clear baseline alterations across lines (**Supplementary Fig. 4B**). Overall, these results indicate that MM proteostasis networks, even if “wired” similarly at baseline, can still drive differential responses under pharmacologic pressure.

### JG98 selectively destabilizes the mitochondrial ribosome

To further explore these unexpected results, we aimed to investigate how the cellular proteome is remodeled after HSP70 inhibition in comparison to blockade of other proteostasis nodes. We performed multiplexed tandem mass tag (TMT) proteomics on three MM cell lines treated with the four compounds above, with each perturbation in biological duplicate (**Fig 4A**). Agents were dosed in each cell line at the LD30 from our drug screens above to ensure cellular drug-responsive phenotype but avoid large-scale cytotoxicity (see Methods). Surprisingly, we found that JG98, but none of the other compounds, led to marked depletion of 55S mitochondrial ribosome subunits when results were aggregated over all three cell lines (**Fig 4B**, **Supplementary Fig. 5A-C**). For JG98, this depletion was highly selective, standing out over essentially all other cellular quantified proteins (4852-5101 protein groups per line, 4033 across all three lines; minimum two unique peptides per protein group) (**Supplementary Fig. 5D-F; Supplementary Dataset 2**). Western blot confirmed marked depletion of the mitoribosome subunit MRPL11 after JG98 treatment but not other drugs (**Supplementary Fig. 6A**). Additionally, the mitochondrial enzyme SOD2 (Superoxide Dismutase 2) was the most increased protein under JG98 **(Supplementary Fig. 5D-E**), further supporting that JG98 causes a mitochondrial stress response in myeloma cells^66,67^.

**Figure 4.**
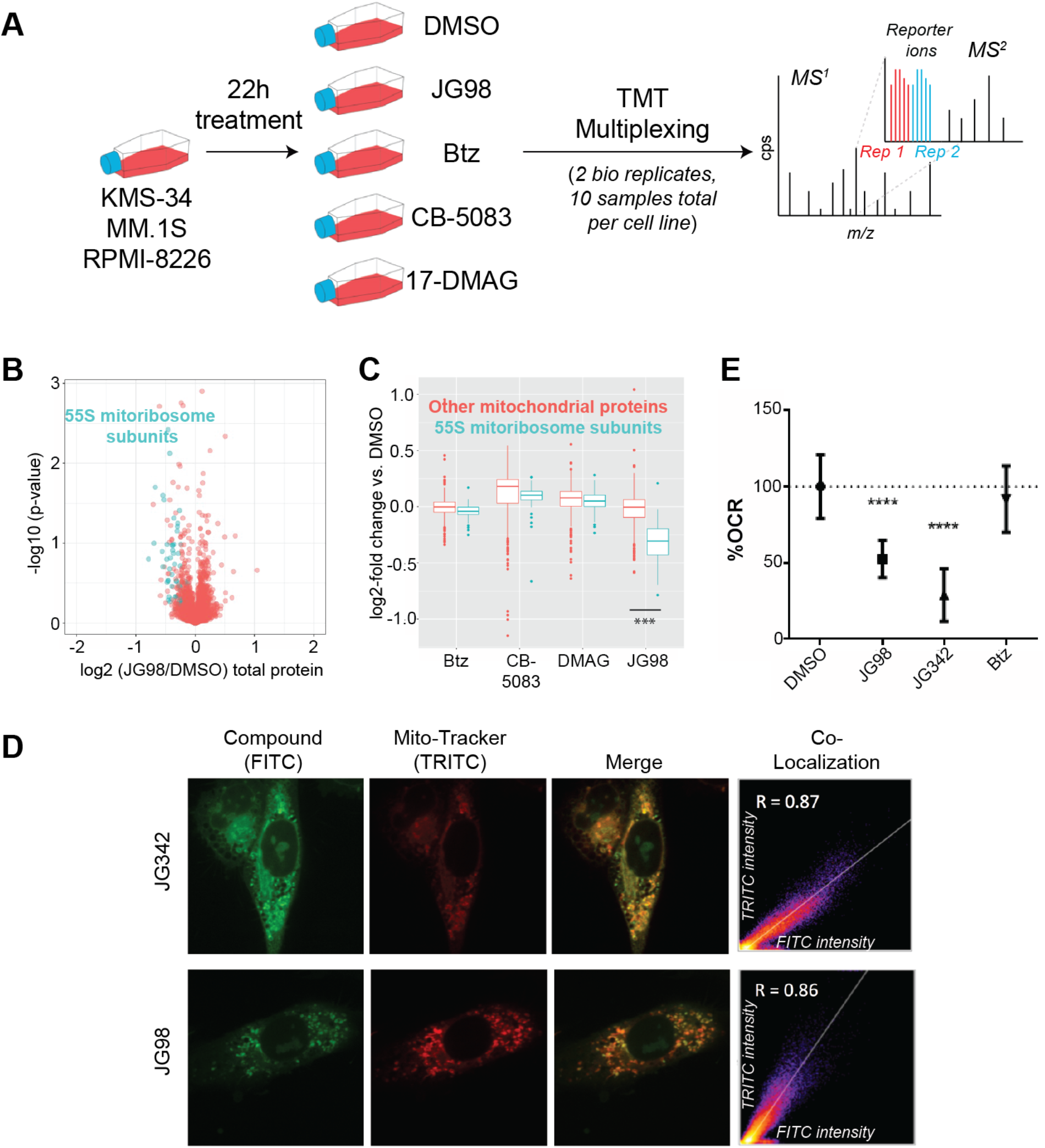
JG98 destabilizes the mitochondrial ribosome. **A.** Experimental schematic. KMS34, MM1.S, and RPMI-8226 MM lines were treated for 22 hr with Bortezomib (Proteasome), 17-DMAG (HSP90), CB-5083 (p97/VCP), JG98 (HSP70), or DMSO. Treatment doses were chosen based on LD30 in drug screens to ensure a cellular phenotype but avoid excessive cell death. (MM1.S cells were treated with 1.75uM JG98, 2.5nM bortezomib, 1uM CB-5083, 150nM DMAG, or DMSO. RPMI-8226 cells were treated with 1.5uM JG98, 7.5nM bortezomib, 1uM CB-5083, 200nM DMAG, or DMSO. KMS34 cells were treated with 2uM JG98, 7.5nM Bortezomib, 1uM CB-5083, 500nM DMAG, or DMSO). Independent biological replicates for each cell line were combined in 10-plex TMT experiments and analyzed by mass spec. **B.** JG98 leads to selective depletion of 55S mitochondrial ribosome subunits; data aggregated across three cell lines (individual cell line data in **Supplementary Fig. 5A-C**). **C.** Log_2_-fold changes for mitochondrial ribosome subunits vs. all other mitochondrial proteins; data aggregated across three cell lines. **D.** JG98 colocalizes with mitochondria in HS5 immortalized bone marrow stromal cells. **E.** JG342 and JG98, but not bortezomib, decrease the oxidative consumption rate (OCR) in HS5 cells. (*n* = 4 wells per treatment, performed 3 times.) *** = two-sided *t*-test *p*-value <0.001; **** = <0.0001.

These findings raised an unexpected hypothesis: as opposed to the predicted effect on cytosolic HSP70’s, could JGs instead overcome PI resistance by inhibiting the mitochondrial HSP70 isoform, mtHsp70/HSPA9/mortalin? Prior work on the precursor molecule MKT-077 suggested that it partitioned into mitochondria but could interact with both mtHsp70 and cytosolic HSPA8^31,32^. To estimate the subcellular localization of JG98, we took advantage of its intrinsic fluorescence^33^. For these live-cell imaging experiments, we used adherent HS5 bone marrow stromal cells, as imaging myeloma cells in suspension at the required resolution was not technically feasible in our hands. After treatment with JG98 or JG342, we visualized mitochondria using a far-red MitoTracker dye to avoid spectral interference from the JG compounds. Indeed, we found that both compounds were almost entirely localized to mitochondria (**Fig. 4D**). As further validation, we performed flow cytometry on intact mitochondria isolated from AMO-1 myeloma cells treated with JG98 and JG342 and found that mitochondria contained high levels of compound (**Supplementary Fig. 6C**). This localization is consistent with other recent data^34^ and might be expected because of the cationic property of JG98 and JG342, as positively charged molecules often localize to this compartment.

To evaluate functional impacts on mitochondria, we performed seahorse respirometry after JG98, JG342, or bortezomib. We found that only the JG compounds led to a decrease in oxygen consumption rate, underscoring a selective mitochondrial perturbation of these agents (**Fig. 4E**). However, we found that neither of these compounds substantially disrupted mitochondrial morphology nor did they lead to mitochondrial depolarization (**Supplementary Fig. 6D-E**). Finally, CRISPR interference-mediated partial knockdown of *HSPA9* in RPMI-8226 myeloma cells stably expressing dCas9-KRAB^35^ (we could not successfully isolate MM cells with CRISPR deletion of *HSPA9*, nor more complete knockdown of *HSPA9*, presumably due to the essential nature of this gene) demonstrated selective loss of MRPL11 but not the cytosolic ribosome subunit RPS6 (**Supplementary Fig. 6F**), illustrating that *HSPA9* knockdown can phenocopy the mitoribosome depletion induced by JG98.

In line with these findings, recent results using two different *in vitro*-generated models of PI resistance also showed that targeting mitochondrial homeostasis may be a selective vulnerability in PI-resistant disease^36,37^ (see also Discussion). Taken together, these results support the notion that JG compounds function primarily via mitochondrial perturbation to drive enhanced efficacy vs. PI-resistant MM.

### Pulsed SILAC illustrates that JG342 leads to loss of nascent mitoribosome subunits

HSP70 family chaperones are primarily thought to function in stabilizing newly-synthesized proteins, though they also may have an array of functions maintaining folded proteins and complexes^38^. We therefore investigated whether the depletion of mitoribosome subunits was primarily due to loss of nascent polypeptides, or, alternatively, destabilization of pre-existing, “mature” proteins.

For these experiments we utilized JG342, to also evaluate if a similar phenotype is present across JG compounds. To distinguish effects on nascent vs. mature proteins and compare multiple timepoints in the same mass spectrometry experiment, we employed a modified version of recently-described methods of TMT combined with pulsed-SILAC proteomics^39, 40^. Most commonly, SILAC (Stable Isotope Labeling of Amino acids in Cell culture) is utilized to compare the steady-state abundance of proteins under two different experimental conditions by incorporating different stable isotope-labeled amino acids resolvable by mass spectrometry^41^. In the design here, though, at *t* = 0 MM.1S cells previously grown in “light” media were switched to “heavy” media containing either JG342 or DMSO as well as stable isotope-labeled arginine and lysine. Three timepoints (16h, 21h, 26h) for both conditions were collected in biological duplicate, and combined into two multiplexed TMT mass spec experiments (**Fig. 5A**).

**Figure 5.**
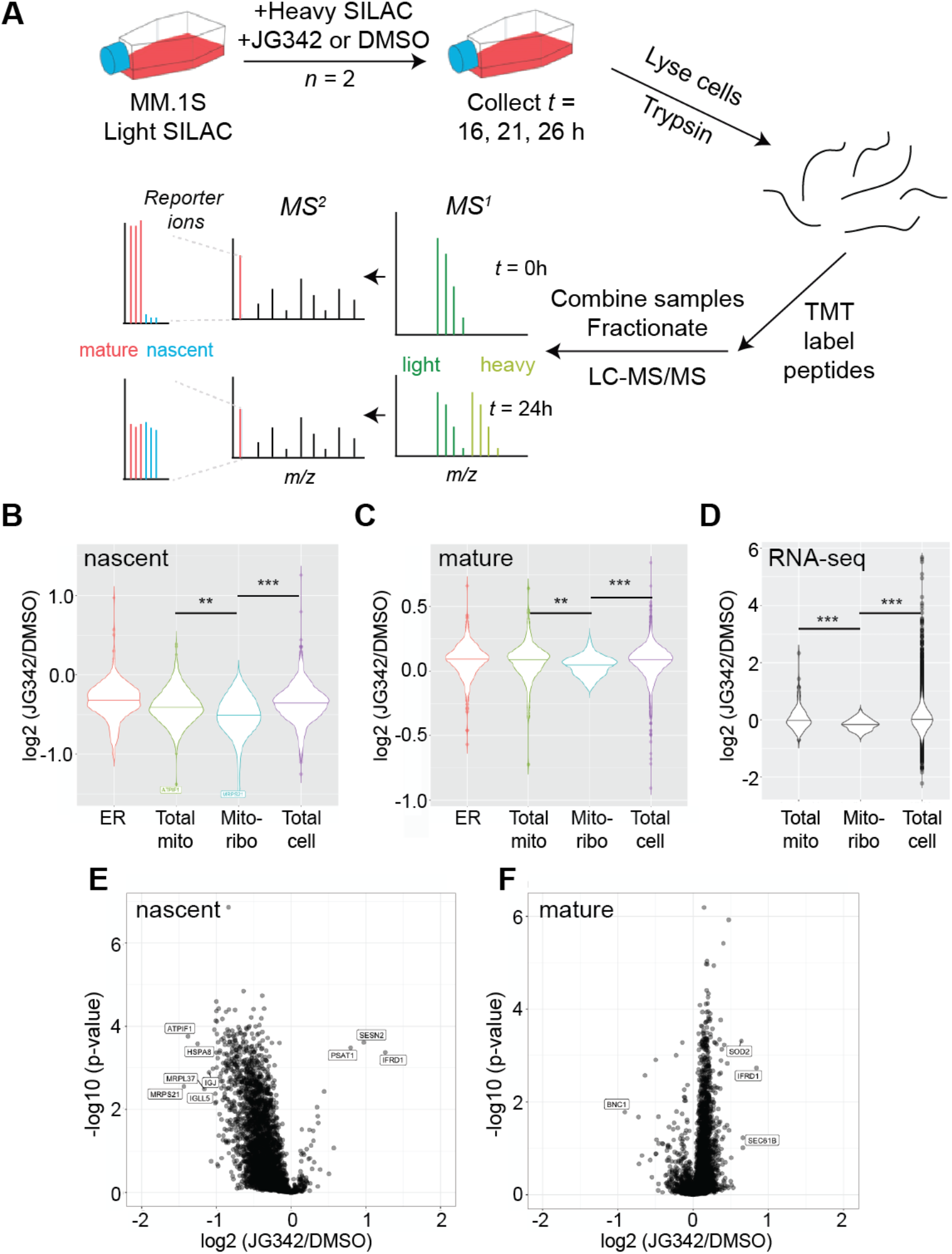
JG342 destabilizes nascent and mature mitochondrial ribosome subunits in the context global translational slowdown. **A.** Pulsed-SILAC experimental schematic. Briefly, MM1.S cells grown in Light SILAC media are switched to Heavy SILAC media containing 350 nM JG342 or DMSO at *t* = 0 hr. Cells are collected at 16, 21, 26 hours, lysed, and proteins digested with trypsin, followed by TMT labeling. Samples are combined in 7-plex experiments, fractionated by HPLC and analyzed by MS/MS on Orbitrap Fusion Lumos. **B-C.** Log_2_-fold changes under 350nM JG342 vs. DMSO across timepoints separating heavy-labeled nascent (**B**) and light-labeled mature (**C**) protein indicate MRPS21 and MRPL37 are among most prominently depleted nascent proteins after JG342. **D-E.** Log_2_-fold changes across timepoints for JG342 treated vs. DMSO for nascent (**D**) and mature (**E**) proteins in endoplasmic reticulum (ER), mitochondria (Total mito), mitochondrial ribosome (Mito-ribo), and rest of the proteome (Total cell, excludes proteins included in previous three categories) demonstrates depletion of mitochondrial ribosome subunits. **F.** RNA-seq TPM Log_2_-fold changes between JG342 and DMSO treated samples from 26 hr timepoint. ** = two-sided t-test p-value < 0.01, *** < 0.001.

In this experiment, monitoring 3104 nascent and 4181 mature proteins (overlap = 2940) (**Supplementary Dataset 3**), we first observed that the large majority of nascent proteins were decreased in abundance after JG342 treatment compared to DMSO control (**Fig. 5B**). This finding was consistent with induction of the Integrated Stress Response (ISR), a conserved response to diverse cellular stresses, where a major outcome is a global decrease in mRNA translation^42^. Consistent with this hypothesis, puromycin incorporation assays biochemically confirmed a global shutdown of protein synthesis (**Fig. 6B**). We also observed marked depletion of nascent 80S ribosome subunits, typically the most prominent translational effect of the ISR^12, 43^ (**Supplementary Fig. 7A-B).**

**Figure 6.**
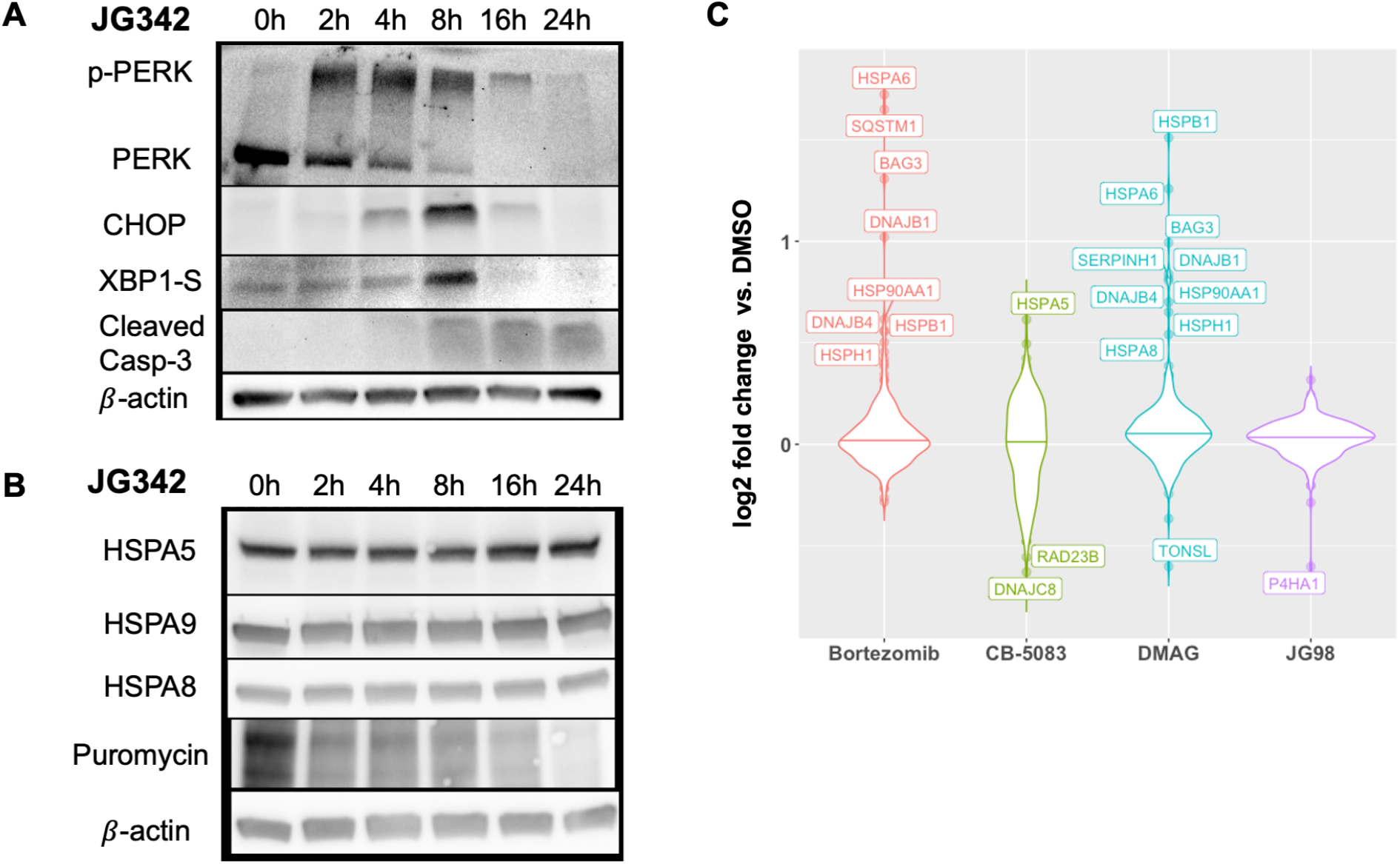
JG compounds activate the UPR without compensatory chaperone upregulation in myeloma cells. **A.** JG342 (3 μM) activates the unfolded protein response in MM1.S cells. **B.** JG342 (2 μM) leads to translational slowdown without upregulation of major HSP70 isoforms HSPA5 (BiP, endoplasmic reticulum), HSPA9 (mitochondria), or HSPA8 (cytosolic). Puromycin incorporation performed by 1 hr incubation of 1μM puromycin at each designated time point. **C.** TMT-proteomics (data from experiment outlined in **Fig 4A**) identifies relative lack of compensatory chaperone upregulation in JG98-treated cells. Analysis here shows 244 proteins out of 441 curated proteostasis genes quantified in TMT mass spectrometry experiments across three cell lines.

However, even among this overall decrease in nascent proteins, we observed that two of the most-depleted proteins were subunits of the mitoribosome (MRPS21 and MRPL37) (**Fig. 5B**). Furthermore, we noted that the overall set of mitoribosome subunits was significantly depleted from the nascent proteome fraction compared to either the general pool of cellular proteins or the specific set of other mitochondrial proteins (**Fig. 5D**). We also we noted decreases, albeit less prominent, in mature mitoribosome subunits (**Fig. 5E**) and mitoribosome mRNA (**Fig. 5F**), suggesting possible effects on the mitoribosome at both the transcriptional and translational levels. Furthermore, SOD2 was among the most enriched mature proteins under JG342, indicative of a mitochondrial stress response^44,45^ (**Fig 5C, Supplementary Fig. 6B).** While the underlying mechanisms remain to be elucidated, these results are consistent with the notion that JG compounds at least partially exert their effects on MM cells through perturbing mitochondrial proteostasis.

### JG compounds induce the UPR and perturb proteostasis without compensatory heat shock response or chaperone upregulation

We previously observed that targeting MM proteostasis with the PI, bortezomib, led to a marked increase in cytosolic chaperone expression, potentially serving as a compensatory response^12^. Here, we asked whether allosteric HSP70 inhibitors might produce similar effects. Prior work in another cancer model showed that allosteric HSP70 inhibition can activate the UPR^46^. In MM cells, we confirmed by Western blotting of canonical markers (spliced XBP1, phosphorylated PERK, CHOP)^47^ that both JG98 and JG342 led to activation of the UPR (**Fig. 6A-B, Supplementary Fig. 8A-C, Supplementary Table 1**). Importantly, we did not observe upregulation of ER, mitochondrial, or cytosolic HSP70 isoforms, nor an increase in master transcriptional regulators of the heat shock response (*HSF1*/*2*/*4*) during treatment with JG98 or JG342 (**Fig. 6B-C; Supplementary Fig. 8A,D**). These proteome- and transcriptome-level findings are in line with prior studies in other systems with Western blotting for selected chaperones^15,16^.

This result with JGs stands in direct contrast to effects of Bortezomib, 17-DMAG, and CB-5083. Despite similar effects on global translation (**Supplementary Fig. 8D**), these other three agents all drove increases in chaperone levels (**Fig. 6C**). Bortezomib and 17-DMAG had partially overlapping response profiles, with increases in small heat shock protein HSPB1, co-chaperones DNAJB1, BAG3, and HSPH1, HSP40 homolog DNAJB4, cytosolic HSP70 chaperone HSPA6, and HSP90 isoform HSP90AA1. In contrast, p97 inhibition with CB-5083 led to a marked increase in the ER-resident HSP70 isoform BiP (HSPA5). Taken together, these results suggest that allosterically inhibiting HSP70 may be an advantageous strategy to target proteostasis, as it does not lead to other compensatory mechanisms to buffer unfolded protein stress.

### Increased proteostasis capacity broadly drives poorer outcomes in myeloma and HSPA9 expression is among the strongest predictors of outcome

Thus far, our collective results suggest that JG compounds are particularly effective versus PI-resistant MM by preferentially targeting the mitochondria-resident HSP70 isoform HSPA9. This finding raises the hypothesis that increased mitochondrial proteostasis capacity, and particularly expression of HSPA9, may also be a feature of PI resistance in MM patients. To investigate this hypothesis, we examined CD138+ tumor cell transcriptomic data from 773 newly-diagnosed patients in the Multiple Myeloma Research Foundation CoMMpass database (IA14 Release) (compass.themmrf.org). The large majority (94.5%) of these patients received a PI as part of their upfront induction therapy.

We first evaluated the possible correlation between overall survival (OS) of patients and the levels of HSP70 isoforms, specifically comparing patients in the top and bottom expression quintiles at diagnosis. Indeed, we found that *HSPA9* expression was by far the strongest predictor of outcome among these genes, where patients in the upper quintile of *HSPA9* expression had a median OS of 58 months, while those in the lowest quintile did not reach median OS (*p* = 2.51e-5) (**Fig. 7A**). In contrast, expression of cytosolic HSP70 isoforms as well as the ER-resident HSP70 *HSPA5* (BiP), led to considerably less pronounced differences in OS (**Fig. 7B-C and Supplementary Fig. 9A-B**). Given that our results suggest a link between HSPA9 inhibition and mitoribosome depletion, we further built an aggregate expression score across all 60 genes comprising the mitoribosome expressed in CoMMpass patients. We found that patients in the upper quintile of mitoribosome expression showed markedly poorer outcomes (*p* = 3.32e-5) (**Fig. 7D**), with a Kaplan-Meier plot closely mimicking that of *HSPA9*.

**Figure 7.**
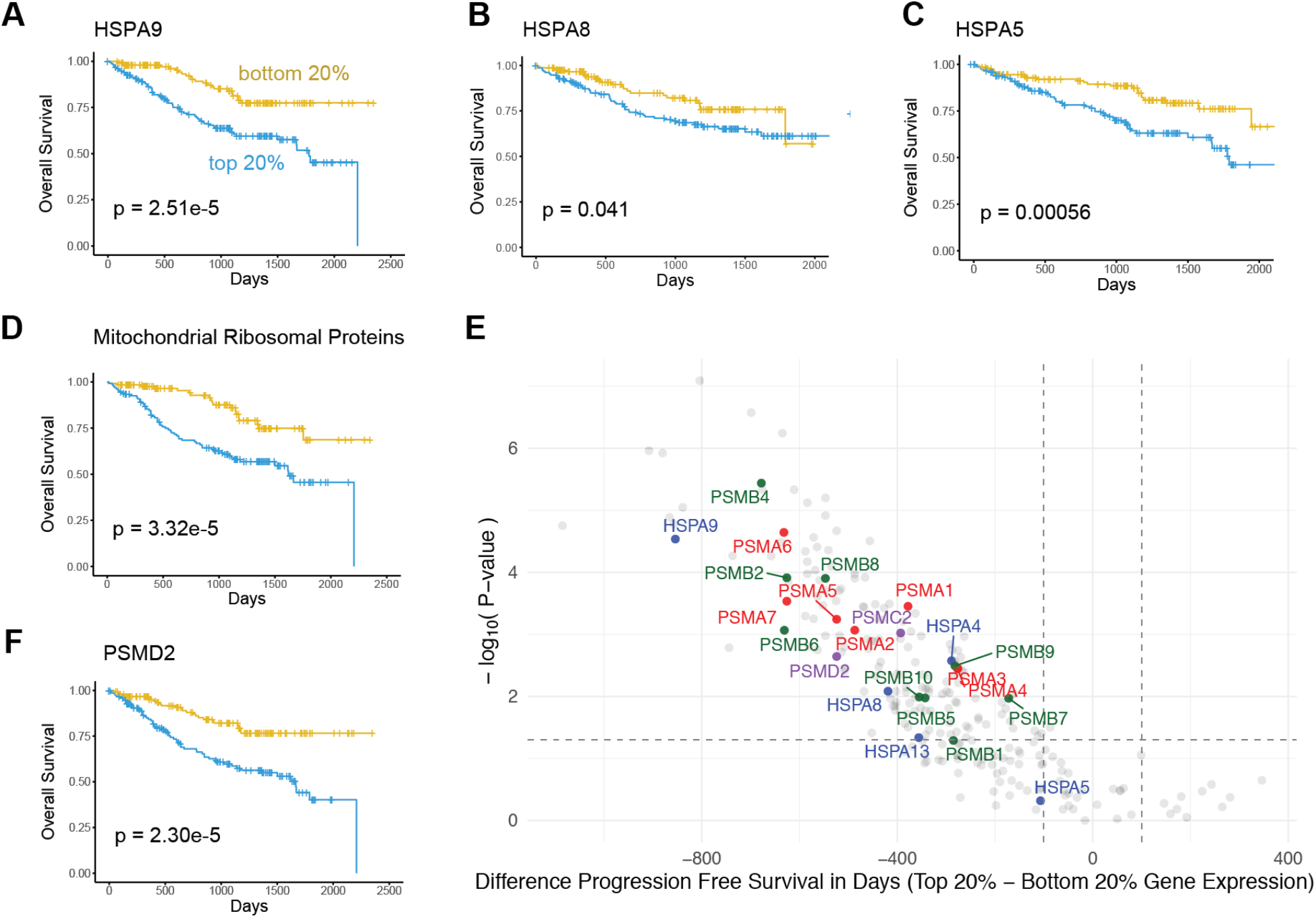
High baseline expression of proteostasis genes, especially HSPA9 and mitoribosome subunits, lead to poorer outcomes in MM patients treated with PIs. **A-D**. Kaplan-Meier curves for overall survival stratified by top and bottom 20 percent of patients for RNA-expression of *HSPA9* (mitochondrial HSP70), *HSPA8* (cytosolic HSP70), *HSPA5* (ER HSP70), and an aggregate 60 gene score of mitoribosome subunits. All RNA-seq (in TPM, from CD138+ enriched tumor cells at MM diagnosis) and overall survival data from 773 patients in the Multiple Myeloma Research Foundation CoMMpass study release IA14. **E.** Volcano plot of proteostasis genes using *p*-values for predictions of relative progression-free survival (PFS) for top and bottom 20% of patients by gene expression (in TPM). 241 genes from 441 from curated proteostasis gene list are included. Genes were included if expressed at TPM >1 across all MM samples and had reached median PFS value in both the top- and bottom-quintiles of expression. Difference in median PFS in days shown along x-axis. *HSPA9* is one of the strongest predictors of poor PFS when highly expressed. **F.** Kaplan-Meier curves for RNA-expression of *PSMD2* shows decreased overall survival in patients with high PSMD2 at baseline. All *p*-values by log-rank test.

While *HSPA9* had the strongest correlation with outcomes, expression of other HSP70 isoforms all showed a similar trend: increased chaperone expression led to shorter survival. This finding raised a general hypothesis for MM plasma cell biology. We speculated that broad increases in tumor proteostasis capacity, potentially through multiple different mechanisms, could decrease sensitivity to PIs, ultimately driving poorer outcomes for patients. To test this hypothesis, we evaluated the difference in median CoMMpass patient Progression-Free Survival (PFS) for the upper and lower expression quintiles across our curated proteostasis gene list in **Fig. 1A** (we note that in this analysis we used PFS instead of OS since this led to more genes with a calculable median survival difference in both the upper and lower quartile of expression). Of these 441 genes, 241 had detectable expression across patients (TPM >1) and were included in our analysis. We found that for the large majority of these genes (160 of 241), increased expression (top quintile) of the proteostasis-related gene led to significantly (*p* < 0.05) poorer survival when compared to low expression (bottom quintile) (**Fig. 7E**). Remarkably, we found no cases where the opposite was true, with bottom quintile gene expression leading to significantly worse outcomes than top quintile.

In this analysis we also note that differential *HSPA9* expression was one of the strongest indicators of patient PFS (**Fig. 7E**). In fact, *HSPA9* was more highly ranked than almost all the 20S core subunits of the proteasome. Taken together, these data underscore the broad ability of MM plasma cells to resist PI treatment if they have greater global capacity to decrease unfolded protein load, potentially mitigating the apoptotic response to unfolded protein stress.

Furthermore, these findings also suggest a possible leading role of HSPA9 and the mitoribosome in maintaining proteostasis in the PI-resistant state, thereby establishing a selective therapeutic vulnerability in these tumors that can be exploited by JG compounds.

### Low 19S cap expression and HSP70 network adaptation do not play a clear role in MM patient PI resistance

Prior analyses using unbiased genetic knockdowns in cell line models^10, 11^ have suggested that low expression of subunits of the 19S proteasome cap may lead to PI resistance. Studies based on this hypothesis have specifically focused on *PSMC2* and *PSMD2*, supported by gene expression data in an older monotherapy clinical trial of bortezomib in myeloma^48^. In our analysis of CoMMpass data, we therefore expected to see decreased expression of 19S cap subunits leading to poorer PFS. However, we found the opposite: increased expression of 19S cap subunits led to poorer outcomes, running in parallel with findings from the 20S proteasomal core (**Fig. 7E-F; Supplementary Fig. 9C**). The reason for this discrepancy is unclear, though it may relate to effects of therapies administered to CoMMpass patients in combination with PI. Regardless, in this real-world data it is apparent that increased proteostasis capacity, including the 19S cap, globally impacts myeloma outcomes under current PI-containing therapeutic regimens.

Furthermore, PI resistance has been linked in cell line models to increased expression of HSP70 family chaperones^7,27^, suggesting potential rewiring of proteostasis to adapt to PI therapy in patients. We therefore obtained data for 50 CoMMpass patients (release IA14) with paired tumor RNA-seq at first relapse and initial diagnoses, all of whom received a PI as part of first-line therapy. Surprisingly, we did not observe any shift toward increased gene expression of any HSP70 isoform, including *HSPA9*, in the relapsed setting (**Supplementary Fig. 9D**). Notably, no increase in expression of proteasomal subunits was observed either (**Supplementary Fig. 9E**). Combined with our survival analysis above, these findings indicate that baseline chaperone expression levels between patients may be more relevant to governing intrinsic resistance to upfront PI therapy, rather than acquired resistance within the same patient after initial response to PI.

## Discussion

Here we aimed to characterize a new approach to address PI resistance, a long-standing issue in MM clinical care. We found that the JG series of allosteric HSP70 inhibitors preferentially eliminated both intrinsic and acquired models of PI resistance. Quantitative proteomics and cellular validation in this setting demonstrated that these compounds function primarily by engaging the mitochondrial-localized HSPA9, rather than through primary effects on the cytosolic HSP70s as expected. While our *in vivo* studies revealed hurdles to progression of these compounds as true clinical candidates, these molecules serve as important chemical probes, elucidating the potential of targeting HSPA9 as means to selectively eliminate PI-resistant disease. This conclusion is supported by analysis of MM patient tumor data, which also revealed the critical role of global proteostasis capacity in MM outcomes.

Overall, our findings fall in line with two recent studies suggesting that mitochondrial homeostasis is a selective vulnerability in PI-resistant disease. In one study, this conclusion was drawn based on bioinformatic analysis of patient tumors and functional analysis of cell lines expressing a low level of the 19S proteasomal cap gene *PSMD2*, demonstrating a strong dependency of PI sensitivity on mitochondrial metabolism^37^. In parallel, a second study involved characterizing *in vitro-*evolved PI-resistant AMO-1 MM cell lines (including the same one we employed here for our initial drug screens (**Fig. 2A-C**)). This recent work demonstrated specific alterations in mitochondrial homeostasis in the PI-resistant state^36^. Targeting PI-resistant AMO1 cells with mitochondrial perturbagens, such as Bcl-2 and AMPK inhibitors, led to increased cell death versus their sensitive counterparts^36^.

Therefore, our results demonstrate that targeting HSPA9 could be a new approach to achieve this same mitochondrial-directed outcome in PI-resistant MM. We do note, though, that multiple lines of prior evidence, as well as our genetic co-dependency analysis here, suggested that cytosolic HSP70s, or perhaps the ER-resident HSP70 HSPA5/BiP, would be good targets in PI-resistant MM^12,49,50^. In other disease models, JG analogs and the precursor MK-077 have been shown to act through cytoplasmic HSP70s^46,51,52^ or to bind both cytoplasmic and mitochondrial HSP70s^32^. Therefore, although these chemical probes inhibit multiple HSP isoforms, the specific biology of the system may dictate that one isoform is more important than the other. For example, despite similar localization data by microscopy, it certainly remains possible that JG342 exhibits overall increased potency versus both PI-resistant and -sensitive MM cell lines compared to JG98 (**Fig. 2C**) by inhibiting additional HSP70s beyond HSPA9. CRISPR interference (CRISPRi) screening of different JG compounds suggest common inhibition of HSPA9 but increased inhibition of HSPA5 in “200” and “300” series compounds (Z. Young and J.E.G., manuscript in preparation). We cannot rule out that molecules strongly targeting only cytosolic HSP70s, or regulators of the heat shock response, such as HSF1, would also have beneficial effects in the same context^50^. Additionally, based on our genetic co-dependency analysis, molecules more prominently targeting cytosolic HSP70 may lead to synergy when co-treated with bortezomib, as opposed to the additivity we observed for JG98 (**Fig. 3**). Overall, the most relevant model may be one where the apoptotic branch of the UPR, triggered via cytosolic HSP70 blockade, ultimately leads to JG-induced death, but additionally targeting HSPA9, with reduced fitness caused by loss of mitoribosomes, specifically leads to the increased sensitivity of PI-resistant cells. Furthermore, broadly targeting HSP70s may confer advantages over other known proteostasis inhibitors, given the lack of compensatory chaperone upregulation after treatment (**Fig. 6C**).

There are limitations to our study. In particular, our experimental results derive from analysis of multiple MM cell line models. We were unable to evaluate differential drug sensitivity in primary myeloma samples due to the fluorescence properties of the JG compounds. Furthermore, the JG scaffold may have significant PK liabilities that complicate further development as a clinical therapeutic. In addition, while our experiments show that JGs robustly localize to mitochondria, lead to mitoribosome depletion, and their effects are pheno-copied by *HSPA9* genetic depletion, we cannot definitively demonstrate that HSPA9 engagement confers this phenotype of increased potency vs. PI-resistant cells. However, the independent observations that PI-resistant cells are highly dependent on mitochondrial metabolism^36,37^, as well as a very recent study confirming that JG treatment pheno-copied mechanistic effects of HSPA9 knockdown in another cancer model^34^, both directly support this notion. Overall, our findings strongly suggest that compounds preferentially targeting HSPA9 will be an exciting approach to overcome PI resistance. These conclusions are supported by our survival analysis of CoMMpass patients.

We can also put these results into context of our initial CRISPR screen data and combination therapeutic analysis. Ultimately, our co-dependency analysis suggests that genes strongly anti-correlated for genetic dependence with proteasome subunits, including HSPA9 (**Fig. 1A**), may prove to be the best targets in the PI-resistant state. In this scenario, cells sensitive to genetic depletion of proteasome subunits at baseline may evolve, upon PI-resistance, to become dependent on proteostasis nodes they previously did not rely upon for survival. This evolved dependence may occur even in the absence of changes in gene or protein expression. Future work will investigate this proposal.

In conclusion, our results support allosterically targeting mitochondrial-localized HSP70 as a promising therapeutic strategy in MM. Our study also reveals global roles of proteostasis networks in the cancer most characterized as highly dependent on this biological process.

## Acknowledgments

We thank Drs. Tom Martin, Nina Shah, Sandy Wong, and Jeffrey Wolf for insightful discussions. We thank the Multiple Myeloma Research Foundation for sponsoring the CoMMpass study. We thank Dr. Martin Kampmann for providing sgRNA sequences for HSPA9 knockdown and CRISPRi cell line. Funding for this study was provided by The Gabrielle’s Angel Foundation for Cancer Research, the UCSF Stephen and Nancy Grand Multiple Myeloma Translational Initiative, NIH DP2 OD022552, K08 CA184116, and R01 CA226851 (to A.P.W.), F32CA236156 (to K.M.T), R01 NS059690 and DoD PC180716 (to J.G.), and UCSF Helen Diller Family Comprehensive Cancer Center P30 CA082103 (to B.H.).

## Author Contributions

I.D.F., C.L., H.S., J.G. and A.P.W. conceived and designed the study. I.D.F., C.L., H.S., M.H., K.M.T., M.C.M., V.S., D.W., P.P., S.T.T., and B.H. performed experiments and analyzed data. Y.-H.T.L. performed bioinformatics analyses. C.D. and B.V.N. provided cell line reagents. B.H., J.G. and A.P.W. obtained funding. I.D.F. and A.P.W. wrote the manuscript with input from all authors.

## METHODS

### DepMap Correlation Analysis

Genetic sensitivity data from genome-wide CRISPR knockout screens for 17,634 genes in 558 cell lines were downloaded from the Cancer Dependency Map (19Q1 Release). Of the 441 manually curated proteostasis genes (see **Supplementary Table 2** for gene names and references), 406 were identified in the CRISPR dataset. Pearson correlation of the sensitivity scores across all cancer cell lines was computed for every pairwise combination of proteostasis genes. The results were displayed as a 406-by-406 heatmap clustered using the complete-linkage method with Euclidean distances. Interactive heatmaps and additional analyses of gene expression and hybrid CRISPR-vs-expression correlations are accessible through a web-based application built using the Shiny package (version 1.4.0.2) in R (https://tony-lin.shinyapps.io/proteostasis-map/).

### Expression-Based Survival Analysis

Survival and gene expression data on CD138^+^ tumors from myeloma patients were downloaded from the CoMMpass database (Build IA14). The expression dataset in transcripts per million (TPM) for 57,996 genes on 908 patient tumors was filtered to retain only newly-diagnosed patient samples (783 total from 773 unique patients identifiers) and genes with TPM > 1 across all samples. Sample duplicates (10 total) were averaged. Next, the overall and progression-free survival data were stratified by expressions in genes of interest, separating the samples by the top and bottom 20% of TPM levels. Kaplan-Meier curves and log-rank P-values indicating statistical difference in survival between the quintiles were generated using the survival and survminer (version 0.4.6) packages in R. For multiple genes (i.e. mitoribosome analysis), a rank order is first assigned on a per-gene basis followed by averaging the ranks across all genes to obtain an aggregate expression score for sample stratification. To summarize the survival differences based on expression levels for many genes, a volcano plot displaying *p*-value versus median difference in survival was generated.

### Differential Expression Analysis

Expression data for 57,996 genes on 908 CD138^+^-enriched myeloma patient samples were downloaded from the CoMMpass database (Build IA14). Using the DESeq2 (ref.^53^) package in R, differential expression analysis was performed on 50 paired tumors with sample collection at initial diagnosis and upon first relapse. The expression data in counts were submitted as input, and sample pairing was indicated in the design formula. The statistical significance of the difference in gene expression between the newly-diagnosed and relapse samples was plotted against expression fold changes in a volcano plot.

### CCLE RNA-sequencing data analysis

Cancer Cell Line Encyclopedia RNA-sequencing data^54^ was downloaded from Cancer Dependency Map (file used was CCLE_depMap_19Q1_TPM.csv). Data from cell lines of interest was extracted for analysis in R.

### Cell culture conditions

Cell lines were maintained in RPMI-1640 medium with 10% FBS (Gemini Benchmark). IL-6 dependent cell lines were cultured in the presence of 50 ng/mL recombinant human IL-6 (ProSpec). Proteasome inhibitor resistant cell lines were cultured in 90nM Bortezomib or Carfilzomib as previously described^9, 27, 55^. Cell lines were authenticated by DNA short tandem repeat profiling at ATCC.

### Monotherapy and combination dose-response viability assays

For monotherapy dose response assays, cells were seeded in 384 well plates using the Multidrop Combi (Thermo Scientific), with 1000 cells seeded in 45 uL media. Drugs were added 24 hours after seeding and viability was measured using Cell-Titer Glo (Promega) 48 hours after the addition of drugs. Combination assays were conducted similarly to monotherapy, with both drugs added 24 hours after seeding and Cell-Titer Glo assay performed 72 hours after addition of drug. Monotherapy dose response assay measurements were performed in quadruplicate and combination assay measurements were performed in triplicate. ZIP synergy scores were processed using SynergyFinder software ^30^.

### Mitochondria isolation

Mitochondria were isolated with the Mitochondria Isolation Kit for Cultured Cells (Thermo, 89874). Briefly, 20e6 MM1.S or AMO1 myeloma cells were treated with 1 uM JG compound or vehicle for 45 minutes prior to addition of MitoTracker Deep Red (Thermo, M22426). After another 45 minutes, cells were harvested, washed with PBS, and 18e6 cells were collected for mitochondria isolation following kit instructions, while 2e6 cells were plated in PBS + 5% FBS for flow cytometry. Flow cytometry was performed on BD Cytoflex (Beckman Coulter). Data was analyzed in FlowJo 8.8.6.

### Mitochondrial membrane potential

Mitochondrial membrane potential was measured by carbocyanine dye DiIC1(5) ((1,1′,3,3,3′,3′-hexamethylindodicarbo-cyanine iodide) staining and flow cytometry as similarly described (^56^, which employs the use of an analogous molecule, the cyanine dye JC-1). Briefly, cells were treated with inhibitors for 24 hours in 96-well plates prior to addition of 50 nM DiIC1(5) (Invitrogen M34151) and 100 uM CCCP (carbonyl cyanide 3-chlorophenylhydrazone, which disrupts mitochondrial membrane potential) to control wells for 30 minutes. Media was removed and cells were resuspended in 5% FBS in D-PBS prior to flow cytometry analysis on BD Cytoflex. Data was analyzed in FlowJo 8.8.6, with a gating strategy including debris and doublet exclusion.

### MitoTracker staining

Mitotracker deep red FM (Invitrogen, M22426) was solubilized in DMSO to yield 100 μM frozen aliquots which were diluted into media to yield a 10 μM stock which was added directly to cell culture media already in the culture yielding a final concentration of 100 nM (to prevent media change derived fluid flow shear stress) 30 min before 4% PFA (MitoTracker red FM was used for PFA fixed samples). Samples were imaged with a Nikon Eclipse Ti spinning disc microscope, Yokogawa CSU-X, Andor Zyla sCMOS, Andor Multi-Port Laser unit, and analyzed with Molecular Devices MetaMorph imaging suite, ImageJ, and ilastik (Gaussian filter).

### Cellular Respirometry

Mitochondrial stress tests were performed with a Seahorse XF24e cellular respirometer on non-permeablized cells at ~80% confluence (50k cells/well) in V7 microplates, with XF assay medium supplemented with 1 mM pyruvate (Gibco), 2 mM glutamine (Gibco), and 5 or 25 mM glucose (Sigma) at pH 7.4 and sequential additions via injection ports of Oligomycin (1 μM final), FCCP (1 μM final), and Antimycin A/Rotenone (1 μM final) during respirometry (concentrated stock solutions solubilized in 100% ethanol (2.5 mM) for mitochondrial stress test compounds). OCR values presented with non-mitochondrial oxygen consumption deducted normalized to DMSO control for clarity.

### Microscopy

Live cell imaging was performed with Nikon Ti-E Microscope equipped with Yokagawa CSU22 spinning disk using the 100X oil objective. Briefly, HS-5 immortalized bone marrow stromal cells were grown overnight in Ibidi 8-well dishes (80826) or MatTek 35 mm dishes (P35G-1.5-7-C). Cells were treated with 500 or 1000 nM JG98 or JG342 and 500 nM Mitotracker Deep Red (Thermo, M22426) for 30 minutes prior to compound washout and imaging. Image processing was performed in ImageJ and Coloc2 was used for co-localization analysis.

### Xenograft Mouse Model

In-house NSG mice were obtained from the UCSF preclinical core facility. 1e6 RPMI-8826-mC/Luc myeloma cells stably expressing luciferase were implanted into each mouse via tail vein injection. Mice were randomized into groups and dosed with 3 mg/kg JG342 three times per week starting on day 18. Tumor burden was assessed by bioluminescence imaging. JG342 was delivered by IV injection at a final concentration of 0.5 mg/mL in 5% DMSO, 10% Cremophor RH40 and Saline. All mouse studies were performed according to UCSF Institutional Animal Care and Use Committee-approved protocols.

### Animal Pharmacokinetics

The animal experiments were carried out in accordance with guidelines of the UCSF Animal Care and Use Committee. Two groups of NSG mice (three each group) were dosed with JG-342 (formulation: 5% DMSO, 10% Cremophor RH 40 and Saline, 0.5 mg/ml) intravenously at 3 mg/kg. Blood samples were collected through tail vein at 5mins, 15 mins, 30 mins, 60 mins, 2h, 6h, 24h, and 48 h time points (4 time points each group). Compound concentrations in plasma were determined by LC/MS/MS, using a previously published protocol^57^.

### HSPA9 knockdown by CRISPRi

Using previously published approaches^58^ we first generated two independent sgRNA’s upstream of the HSPA9 transcriptional start site, designed to repress transcription by the CRISPR intereference dCas9-KRAB fusion protein:

g261l (2162 HSPA9_i1 HSPA9_-_137911022.23-P1P2): GTATCATGGCGGATAAATGG
g314l (2163 HSPA9_i2 HSPA9_-_137911079.23-P1P2): GGAGCTGCGCGATGCGGTGG

To generate lentivirus for stable transduction of these sgRNAs, 2.35 mL of incubated DMEM were aliquoted into four separate wells of a 6-well plate. 9e5 LentiX cells were added to each of these wells. The plate was allowed to incubate overnight at 37oC and 5% CO2.

After 24 hours, the following were added to 4 separate 1.5 mL Eppendorf tubes: 1.35 mg of pCMV plasmid, 0.165 mg of pMD plasmid, and 12 mL of Transfection 5 Reagent. 1.5 mg aliquots of g261l plasmid were added to two Eppendorf tubes, and 1.5 mg aliquots of g314l were added to the remaining two Eppendorf tubes. All four Eppendorf tubes were incubated at room temperature for 15 minutes. The contents of each Eppendorf tube were added to a LentiX cell-containing well of the 6 well plate prepared the prior day. The plate was wrapped in parafilm and was incubated at 37oC and 5% CO2 for approximately 72 hours.

After 72 hours of incubation, a 0.45mM filter was attached to a 3 mL syringe. The supernatant of one well from the six well plate was drawn up through the filter into the 3 mL syringe. The contents of the syringe was carefully ejected into a 15 mL Falcon tube. The tube was parafilmed and frozen at −80oC until future use. The same process was repeated for each well of LentiX cells in the 6-well plate.

For knockdown we utilized an RPMI-8226 multiple myeloma cell line (RPMI-8226 JTC) engineered to stably express the dCas9-KRAB CRISPRi construct at the *AAVS* safe harbor locus^35^. This “insulator” approach avoids CRISPRi construct silencing during cell culture and passage. 1.5e6 RPMI8226 JTC cells were counted and aliquoted into two wells of a 6-well plate. 6 uL of polybrene was added to each well. 1 mL of Lenti virus generated from g261l plasmid was added to one well and 1 mL of Lenti virus generated from g314l plasmid was added to other well. The plate was covered in parafilm and was spinfected for 2 hours at 1000 rcf. The cells were then incubated overnight at 37oC and 5% CO2.

After 24 hours of incubation, the cells from each well were separately washed twice with PBS and resuspended in 10 mL of RPMI media (20% FBS/1% P/S). 2 ug of puromycin were added to each well. After two days, the cells media was changed and an additional 2 ug of puromycin were added to each well. Percentage of BFP positive cells were monitored after each round of selection using the PB450-A channel. After two cycles of puromycin selection, 1e6 cells were spun down and prepared for Western blotting as described above, with comparison to RPMI-8226 JTC cells transduced with scrambled sgRNA ^35^.

### RNA-sequencing and data analysis

RNA extraction, library preparation, and sequencing were performed at BGI (Shenzhen, China). Samples were sequenced using the BGISEQ-500 platform, with an average of 25.75 million reads per sample. For genome mapping, clean reads were mapped to reference genome using HISAT^59^ with an average of 94.2% mapping across samples. For gene expression analysis, clean reads were mapped to reference transcripts using Bowtie2 (ref.^60^) and expression levels calculated using RSEM^61^. Transcripts per million (TPM) were used for data analysis performed in R. Raw sequencing data has been deposited at the Gene Expression Omnibus (GEO) (accession number GSE148659).

### Western blot sample preparation and analysis

For unfolded protein response and chaperone regulation blots, MM1.S cells were seeded at a density of 1e6/mL and treated with 2 or 3 μM JG342 or 3 μM JG98 for 2, 4, 8, 16, or 24 hours as indicated in figure legends. Puromycin treatment at 1 μM was performed at 37 C for 1 hour prior to collection. Cells were collected and washed with PBS, flash frozen, and stored at −80C prior to lysis and western blot analysis. For Puromycin incorporation western blot, 5e6 KMS34 cells were plated in 5mLs of media in 6-well plates and treated for 22 hours with 2μM JG98, 7.5nM Bortezomib, 1μM CB-5083, 500nM DMAG, or DMSO. At 21 hours, cells were treated with 1μM Puromycin (Thermo A1113803) for 1 hour prior to harvest, PBS wash, and storage at −80C. Cells pellets were lysed in 1X RIPA lysis buffer (Millipore) with HALT protease and phosphatase inhibitors (Thermo 78430), incubated on ice for 15 minutes, sonicated for 15 seconds at 1 Hz cycles on ice, and cleared by centrifugation at 17000xg for 10 minutes at 4C. Western blots were performed as previously described^62^. Primary antibodies for immunoblotting were obtained from Cell Signaling (α-PERK 5683S, α-CHOP 2895S, α-Cleaved Caspase 3 9644T, α-B-actin 3700S, α-HSPA5 3183S, α-HSPA9 3593S, α-HSPA8 8444S, α-MRPL11 2066S, α-SOD2 13141T, α-RPS6 5G10), Kerafast (α-Puromycin EQ001), and Biolegend (α-XBP1-S 143F) and used at manufacturer recommended dilutions. Ponceau-S (Thermo BP10310) and total protein stain or anti-β-actin were used as loading controls for immunoblots as indicated in figure legends.

### Drug treatments for mass spectrometry samples

For experiments outlined in Figure 4A, 5e6 cells were seeded in 5 mL media in 6-well plates and treated with compounds for 22 hours at the following concentrations: MM1.S cells were treated with 1.75 μM JG98, 2.5 nM bortezomib, 1 μM CB-5083, 150 nM DMAG, or DMSO. RPMI-8226 cells were treated with 1.5 μM JG98, 7.5 nM bortezomib, 1 μM CB-5083, 200 nM DMAG, or DMSO. KMS34 cells were treated with 2 μM JG98, 7.5 nM Bortezomib, 1 μM CB-5083, 500 nM DMAG, or DMSO. Cells were harvested and washed with PBS, snap frozen in liquid nitrogen, and stored at −80C. For pulsed-SILAC experiments, MM1.S cells were grown in Light SILAC media for at least six passages to allow complete labeling and adaptation to dialyzed FBS. At time = 0, cells were exchanged to Heavy SILAC media with 350nM JG342 or DMSO. Cells were collected at 16, 21, and 26 time points, washed with PBS, snap frozen, and stored at −80C. Light and Heavy SILAC media were composed as follows: Thermo RPMI media for SILAC (88365) supplemented with 10% dialyzed FBS, 70 mg/L Lysine (Sigma) or L-Lysine-^13^C_6_,^15^N_2_ (Cambridge Isotope), 40mg/L Arginine (Sigma) or L-Arginine-^13^C_6_,^15^N_4_ (Cambridge Isotope), 200mg/L Proline (Sigma), and 1% PenStrep.

### Mass Spectrometry sample preparation

Cell pellets were lysed in 6M GdnHCL, 0.1M Tris pH 8.5, with 5mM TCEP and 10mM 2-chloro acetamide. Lysates were sonicated for 45 seconds at 1 Hz cycles on ice and cleared by centrifugation at 16000g for 10 minutes at 4C. Protein concentration was measured with 660 assay (Pierce 22660) and 100 μg protein was digested with 2 μg Trypsin (Pierce 90057) for 18-22 hours with digestion halted by addition of TFA to 1% vol/vol. Acidified samples were centrifuged at 17,200g for 5 minutes at RT to remove precipitate. Cleared samples were desalted on SOLA columns (Thermo 60109001) according to manufacturer instructions and eluted in 50% Acetonitrile with 0.1% FA and vacuum dried prior to storage at −80C. Miniaturized TMT labeling was performed based on modified protocol^63^. Briefly, peptide concentrations were measured using the Pierce Peptide Colormetric Assay (Pierce 23275). 20 ug peptides were resuspended in 17.5 uL 50 mM HEPES pH 8.5 and labeled with 50 μg TMT reagent dissolved in 2.5 μL Acetonitrile for 1 hour at 25C and 500 rpm. Reactions were quenched by adding hydroxylamine to final concentration of 0.4% and incubation for 15 min at 500 rpm. TMT labeled samples were combined, acidified by addition of 0.1% TFA, vacuum dried, and stored at −80C.

### High pH fractionations

For 7-plex pSILAC-TMT, peptides were subjected to high pH fractionation using an XBridge C18 column (1.0×100mm, 3.5um, Waters) on a Waters 2796 Bioseparations Module HPLC machine with non-linear gradient designed as follows: start at 90% Buffer A (5% Acetonitrile, 10mM TEAB) and 10% Buffer B (90% Acetonitrile, 10mM TEAB), ramp to 15% B over 8 minutes, ramp to 27.7% B over 22 minutes, ramp to 46.6%B over 14 minutes, ramp to 55% B over 4 minutes, ramp 90% B over 6 minutes for column wash prior to re-equilibration. Fractions were collected every 30 seconds and every 10^th^ fraction was combined for a final 10 fractions for mass spectrometry analysis. For 10-plex TMT shotgun experiments, peptides were fractionated using a High pH Reversed-Phase Fractionation kit (Pierce, 84868). Briefly, columns were prepared with two acetonitrile washes, followed by two 0.1% TFA washes. Samples were loaded onto the column and washed with HPLC grade water. Peptides were eluted in 8 fractions – 10%, 12.5%, 17.5%, 20%, 22.5%, 25%, and 50% acetonitrile, 0.1% triethylamine. Samples were then vacuum dried and resuspended in 2% Acetonitrile, 0.1% formic acid for mass spec analysis.

### LC-MS/MS operation

For 7-plex pulsed SILAC-TMT LC-MS/MS, 500 ng of peptides were injected into Easy-Spray reversed phase column (Thermo ES800) on a nanoACQUITY UPLC (Waters) coupled to a Fusion Lumos Mass Spectrometer (Thermo) with the following non-linear gradient in which A is 0.1% Formic Acid and B is Acetonitrile plus 0.1% Formic Acid: 8% B to 30% B for 110 minutes, 30% B to 50% B for 20 minutes, 50% B to 70% B for 5 minutes, 70% B to 80% B for 1 minute, and 8% B for 7 minutes to re-equilibrate. For MS1 data acquisition, scan range was set to 375-1500 m/z, AGC target was set to 4e5, and maximum injection time (IT) was set to 50ms. For MS2 data acquisition, isolation window was set to 0.7 m/z, with HCD energy set to 38 percent, orbitrap resolution was set to 50000, and AGC target was set to 1.0e5.

For 10-plex TMT shotgun experiments, 1 μg of peptides were injected into a Dionex Ultimate 3000 NanoRSLC instrument with 15-cm Acclaim PEPMAP C18 (Thermo, 164534) reverse phase column coupled to a Thermo Q Exactive Plus mass spectrometer. HPLC non-linear gradient was as follows with buffer A 0.1% FA and buffer B 0.1% FA in 80% Acetonitrile: 3-8% B for 11 minutes, 8-34% B for 80 minutes, 34-50% B for 15 minutes, 50-70% B for 5 minutes with hold at 70% for 3 minutes, and 99% B column wash for 4 minutes prior to re-equilibration for 13 minutes. For MS1 acquisition, spectra were collected in data dependent top 15 method with full MS scan resolution of 70,000, AGC target was set to 3e6, and maximum IT set to 50ms. For MS2 acquisition, resolution was set to 35,000, AGC set to 1e5, and maximum IT to 100ms with Normalized Collison energy of 32.

### Proteomic data analysis and quantification

Mass spectrometry data was processed in Maxquant^64^ version 1.6.2.1 with the following settings: PSM/Protein FDR were set to 0.01, Carbidomethylation was set as fixed modification and methionine oxidation and N-terminal acetylation were set as variable modifications, minimum peptide length = 7, matching time window set to 0.7 min, alignment time window set to 20 min, and match between runs was used, along with other default settings. Data was searched against the Uniprot Swiss-Prot human proteome (ID:9606, downloaded from Uniprot in 2018). For TMT-pSILAC multiplexing analysis, separate parameter groups were used for heavy and light labeled protein analysis run in the same MS2 experiment using the isobaric labels function in Maxquant. For the heavy parameter group, heavy arginine was set as fixed modification and heavy lysine modifications were added into TMT-tag masses without altering the diagnostic peaks, which denote the TMT cleaved label masses. Proteingroups files were exported from Maxquant, filtered to remove contaminants, and filtered for proteins with at least two unique peptides for analysis. For TMT-pSILAC data analysis the razor peptide was included for unique peptide count threshold. Data analysis was performed in Perseus^65^ and R. Subcellular compartment gene lists were downloaded from Uniprot and restricted to reviewed entries. The mass spectrometry proteomics data have been deposited to the ProteomeXchange Consortium via the PRIDE^66^ partner repository with dataset identifiers PXD018617 and PXD018387.

### Gene Ontology (GO) Analysis

Panther Overrepresentation Tests (Release 20190711) were performed on RNA-sequencing and proteomic datasets. Gene Ontology database used was released (2019-10-09), and FISHER test with FDR correction were used. For RNA-sequencing data, only genes with at least one measured TPM greater than 5 and no zero TPM values were used for background (10212 genes in background list). From this subset, genes with log_2_ fold changes greater than or equal to 1 in both replicates were considered as significantly upregulated and included in the analysis list. For downregulated nascent proteins from TMT-pSILAC experiment, nascent proteins with Z-score less than or equal to −2 and p-value less than or equal to 0.05 (*n* = 883) were used with background list all identified nascent proteins with minimum 2 unique peptides (*n* = 3104).

### Gene Set Enrichment Analysis (GSEA)

GSEA software^67^ was downloaded from https://www.gsea-msigdb.org/. Median normalized TMT mass spec data across three cell lines (experiment outline in 4a) was used for input (4033 genes). Gene set used was h.all.v6.2.symbols.gmt.

## SUPPLEMENTARY FIGURES and TABLE

**Supplementary Figure 1.**
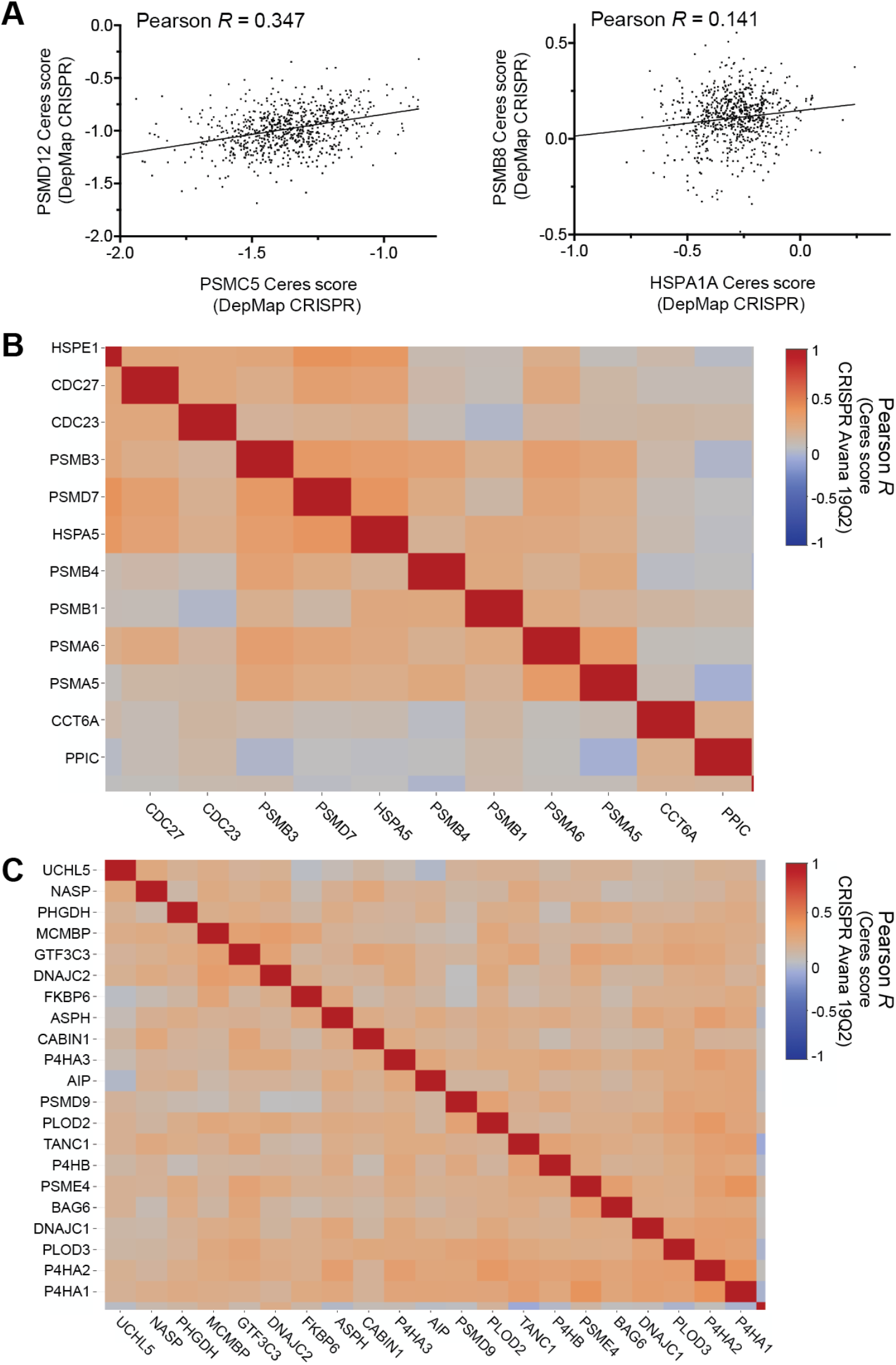
Additional genetic co-dependency clusters. **A.** Crispr-knockout co-dependency validations of gene pairs included in prominent cluster shown in **Fig. 1B. B.-C.** Additional co-dependency clusters as noted in white highlight in **Fig. 1B**. Data from DepMap CRISPR Avana 19Q1 release.

**Supplementary Figure 2.**
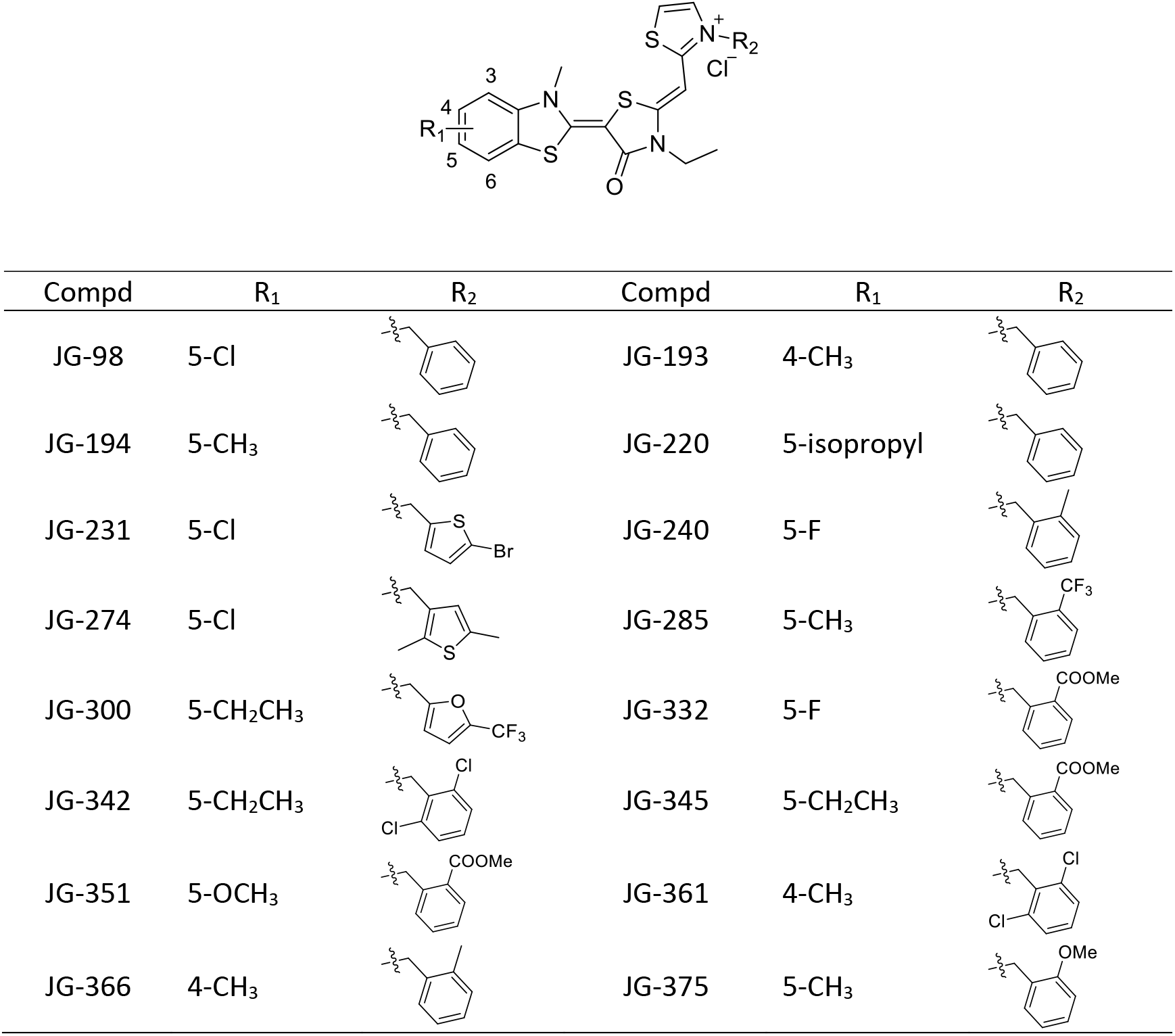
Structure of JG compounds tested in AMO-1 and AMO1-BtzR cell lines. The synthesis and characterization of JG compounds was previously described in ref.^16^.

**Supplementary Figure 3.**
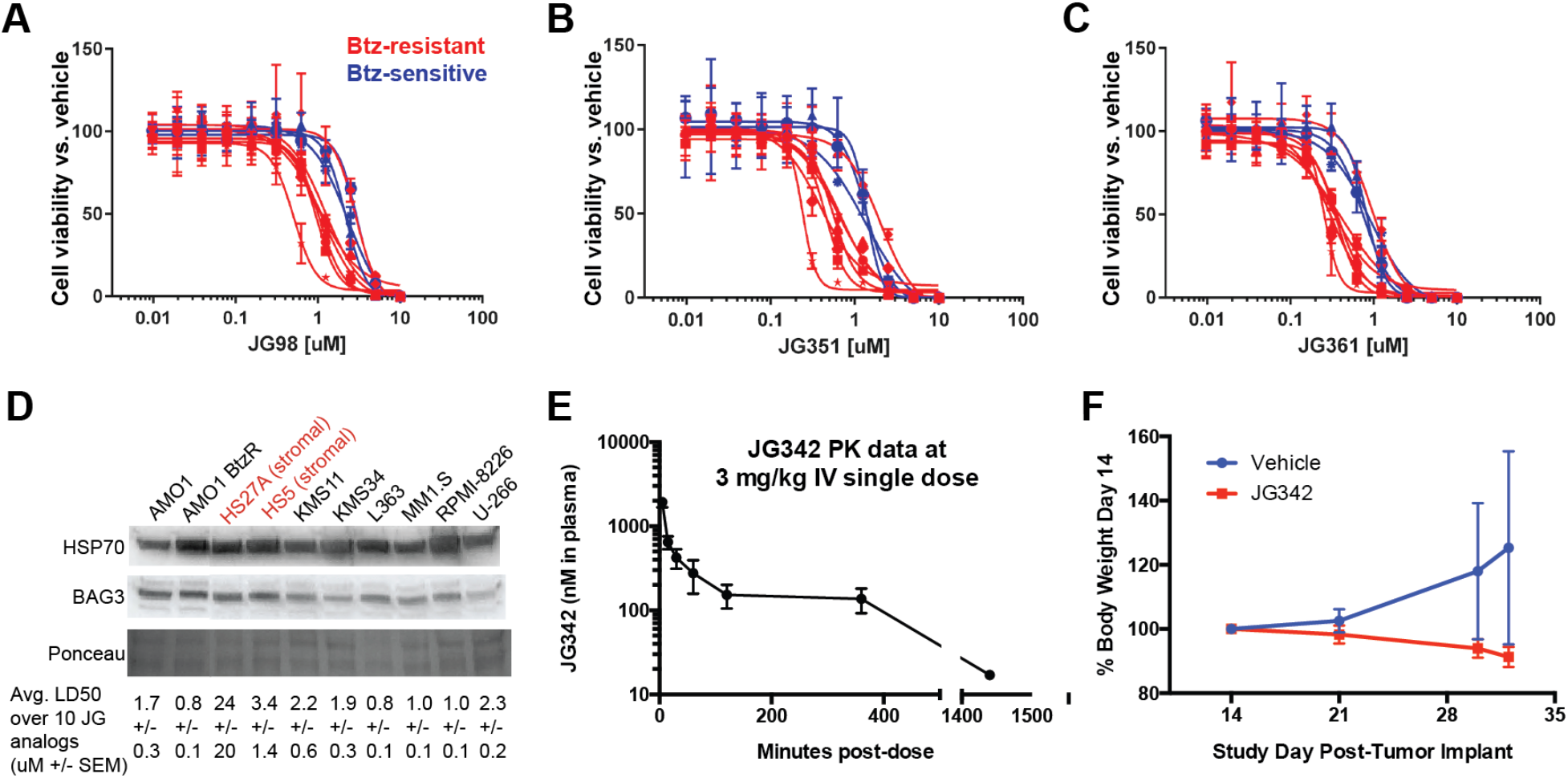
Additional data on JG differential sensitivity and murine studies. **A-C.** Intrinsically and evolved proteasome-inhibitor resistant cell lines are more sensitive to JG98, JG351, and JG361 than intrinsically sensitive lines (same lines used as in **Fig. 2E-G**). **D.** Total HSP70 and BAG3 levels in multiple myeloma cell lines compared with average μM LD50s over 10 JG analogs (LD50 data from **Fig. 2C**). Note that bortezomib-resistant (BtzR) AMO-1 line demonstrates higher expression of total HSP70 than parental AMO-1, consistent with proteomic results of ref.^9^. However, across lines there is no correlation with HSP70 or BAG3 expression and sensitivity to JG compounds. **E.** Two groups of NSG mice (*n*=3 per group) were dosed with JG342 (formulation: 5% DMSO, 10% Cremophor RH 40 and Saline, 0.5 mg/ml) intravenously at 3 mg/kg. Blood samples were collected through tail vein at 5 mins, 15 mins, 30 mins, 60 mins, 2h, 6h, 24h and 48 h time points, and compound concentrations in plasma were determined by LC/MS/MS. **F.** Body weights of mice relative to start of trial for mice treated with vehicle or 3mg/kg JG342 demonstrate limited tolerability of ongoing JG342 treatment (*n* = 5 per arm; data shown +/− S.D.), mouse study as in **Fig. 2H-I**.

**Supplementary Figure 4.**
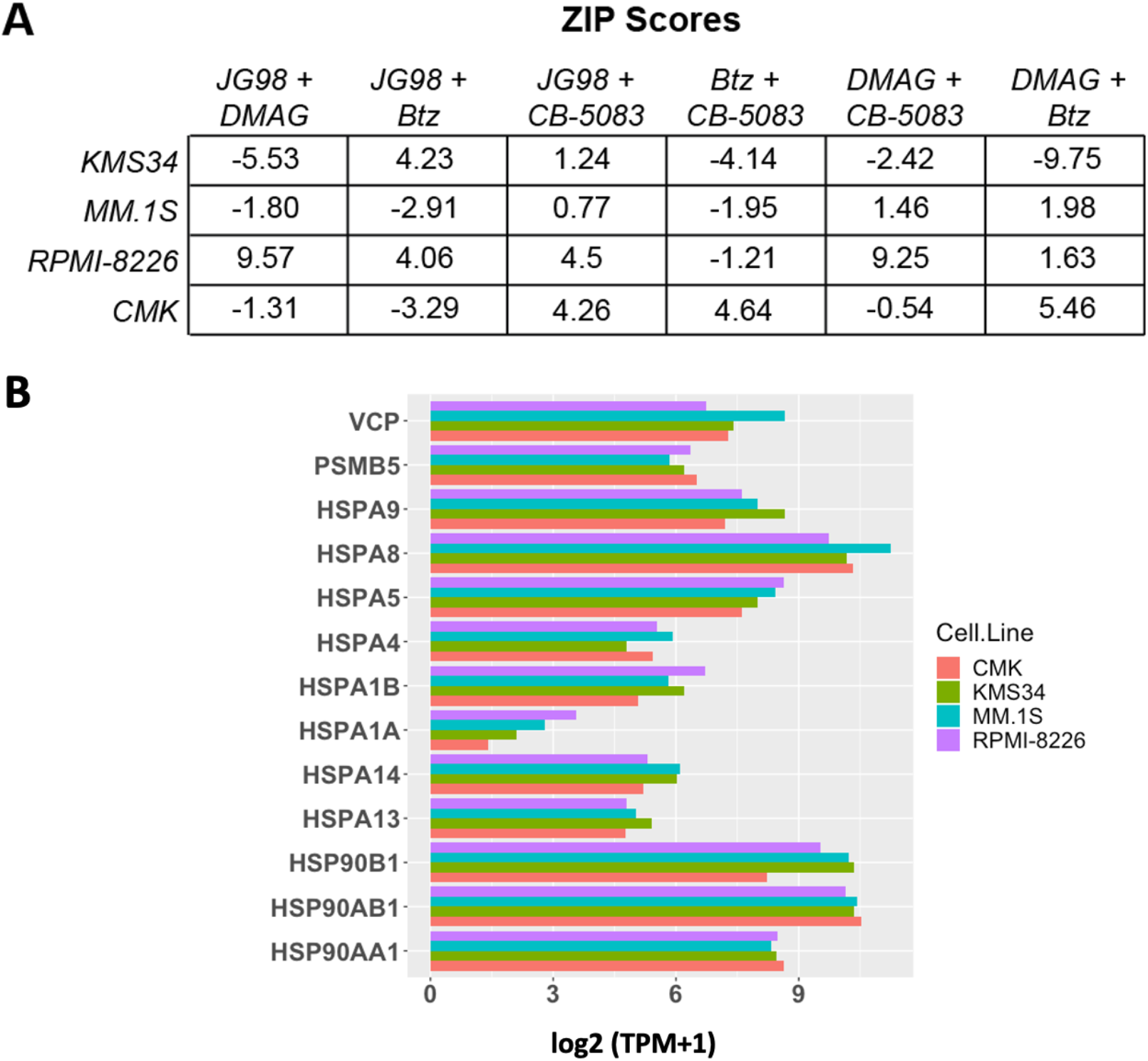
Differential drug combination effects are not influenced by baseline target gene expression. **A.** ZIP synergy scores for 72-hour combination assays in 3 myeloma (KMS34, MM.1S, RPMI-8226) and one AML cell line (CMK), as shown graphically in **Fig. 3C**. **B.** Gene expression from RNA-sequencing (in Log_2_(TPM+1)) for canonical drug targets of 17-DMAG, JG98, CB-5083, and Bortezomib (HSP90 isoforms, major HSP70 isoforms, *VCP*, and *PSMB5*, respectively), shows similar baseline proteostasis gene expression across lines. CCLE gene expression data downloaded from DepMap portal (release 19q1).

**Supplementary Figure 5.**
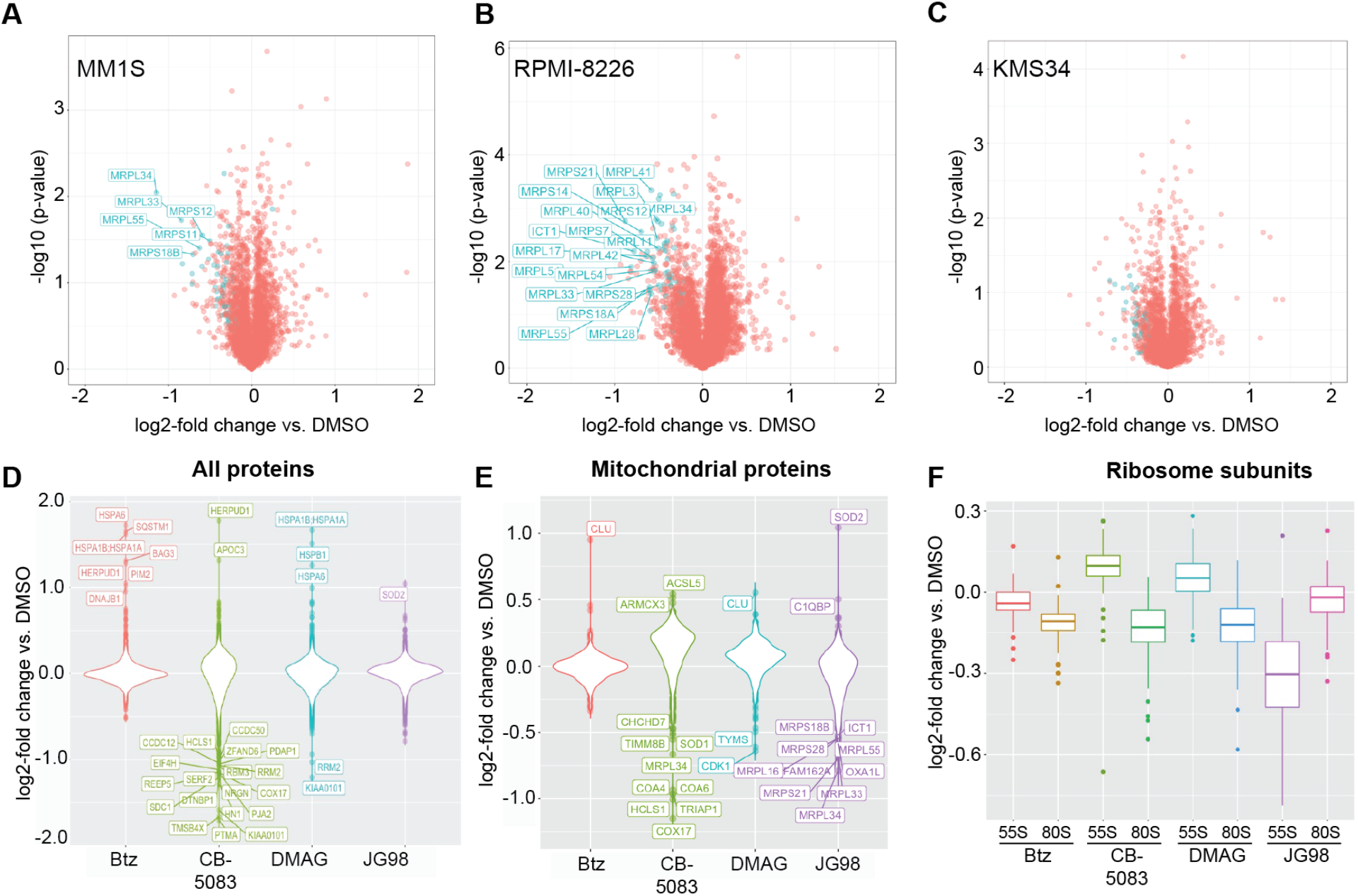
Further proteomic analysis of mitoribosome depletion after JG98 treatment. **A.-C.** Shotgun proteomic data across the three individual MM cell lines included in the aggregate data of **Fig. 4B** demonstrate consistent depletion of mitoribosome subunits (in blue) after JG98 treatment. Labeled mitoribosome subunits have Log_2_-fold change less than −0.5 and *p*-value less than 0.05. **D.-E.** Log_2_-fold changes for whole proteome and mitochondrial proteins after treatment with Bortezomib, CB-5083, DMAG, and JG98, as in **Fig. 4**; data consolidated across three cell lines. **F.** Log_2_-fold changes for drug treatment vs. DMSO for 55S mitoribosome and 80S ribosome subunits, as in **Fig. 4**; data consolidated across three cell lines. Analysis shows that only JG98 has selective depletion of mitoribosome subunits. Box and whiskers plot shows median, 25^th^ and 75^th^ percentile (box) and tails of distribution (whiskers).

**Supplementary Figure 6.**
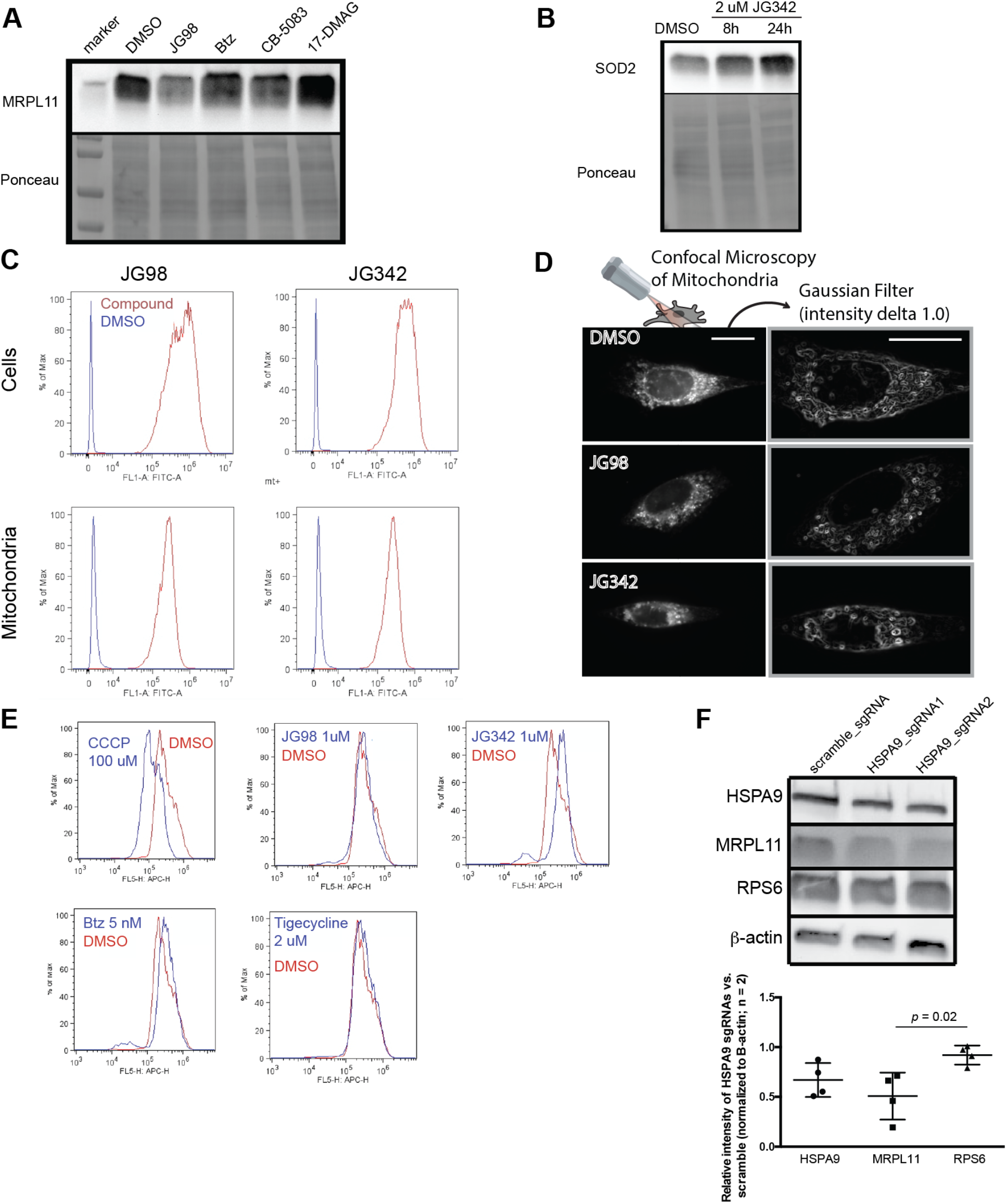
Mitochondrial morphology and mitoribosome depletion validation experiments. **A.** MM.1S cells were treated for 24 hours with 1.75μM JG98, 2.5 nM Bortezomib, 1μM CB-5083, 150 nM DMAG (LD30 doses from drug screens in MM.1S, as used in shotgun proteomics experiment) or DMSO. MRPL11 levels detected by western blot. Ponceau stain shown as loading control. **B.** MM1.S cells were treated with JG342 at 2μM for 8 and 24 hours and SOD2 expression was measured by western blot, confirming findings by proteomics. **C.** AMO-1 cells were treated with 1μM JG342 or JG98 prior to mitochondrial isolation and flow cytometry of isolated mitochondria and intact cells. JG compounds were visualized by FITC fluorescence. **D.** Representative confocal microscopy images of HS5 bone marrow stromal cells treated with 1 μM JG98, 1 μM JG342, or DMSO for 15 hr with 30 min incubation with 100 nM Mitotracker Deep Red prior to fixation (4% PFA, 15 min at 22 ºC) and imaging. **E.** Mitochondrial membrane potential measured by DiIC1(5) fluorescence in MM1.S cells treated with 1 μM JG compounds, 5 nM bortezomib, or 2 μM tigecycline (mitochondrial protein synthesis inhibitor) for 24 hours and analyzed by flow cytometry. 100 μM Carbonyl cyanide chlorophenyl hydrazone (CCCP) is used as a positive control compound known to decrease mitochondrial membrane potential. **F.** RPMI-8226 MM cells with stable integration of the CRISPR interference dCas9-KRAB construct^35^ were transduced with a non-targeting guide RNA or two independent guide RNAs targeting *HSPA9*. Western blotting was performed for HSPA9, MRPL11, and RPS6. Quantification versus beta-actin loading control and with relative fold-change of two *HSPA9* sgRNAs vs. scramble sgRNA, in two independent replicates, demonstrates significant depletion of MRPL11 but not RPS6. *p*-val from two tailed *t*-test.

**Supplementary Figure 7.**
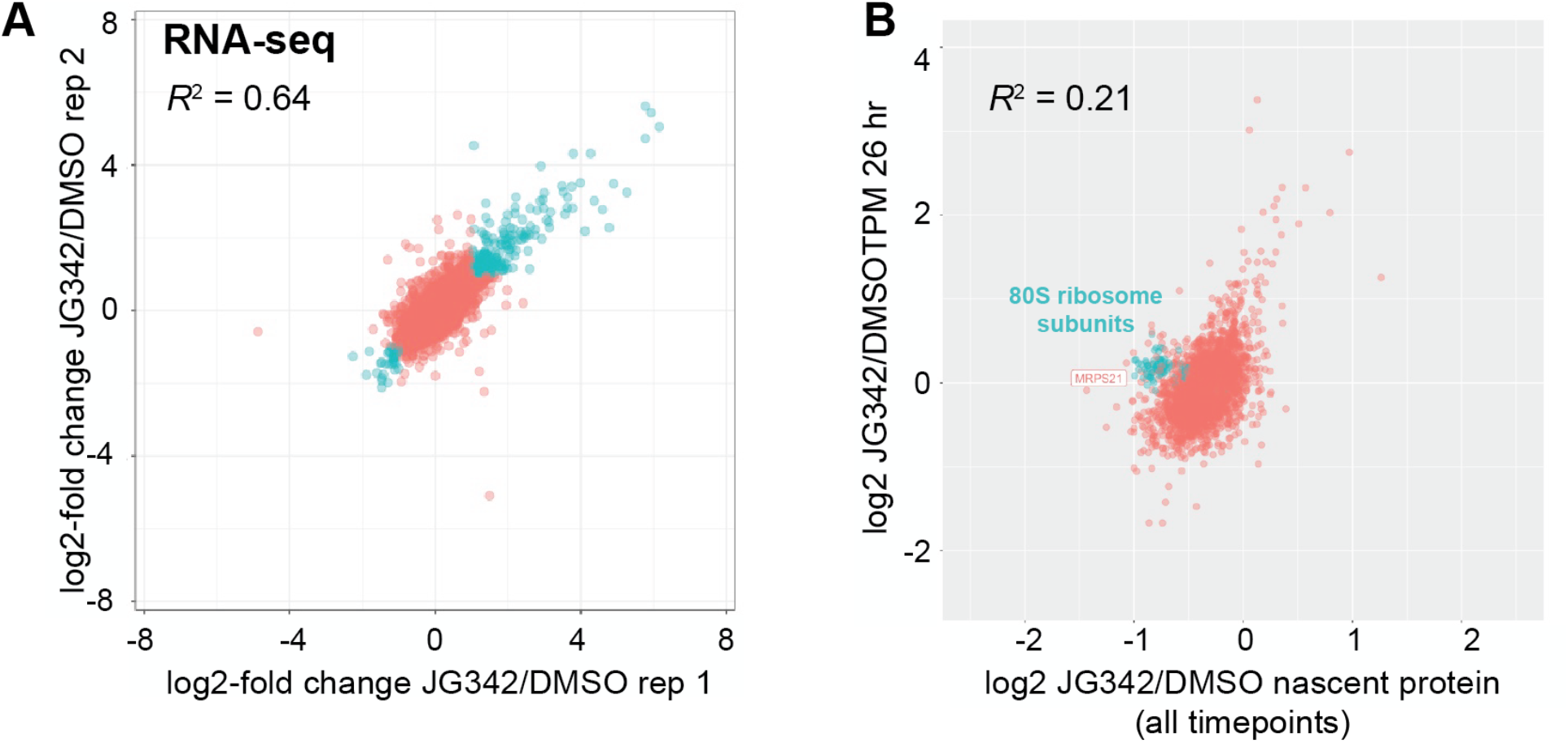
Dynamic transcriptome-proteome alterations after JG analogs. **A.** RNA-sequencing data from 26 hr time points across biological replicates. Genes with Log_2_-fold changes >|1| in both replicates (in blue) were considered upregulated or downregulated. Upregulated genes were used for Gene Ontology analysis in **Supplementary Table 1**. **B.** Log_2_-fold changes for 26 hr time point for RNA and nascent protein across timepoints. 80S ribosome subunits are enriched in post-transcriptionally regulated proteins, consistent with known translational response to drug-induced stress. Mitoribosome subunit MRPS21 is the most-depleted nascent protein among all quantified.

**Supplementary Figure 8.**
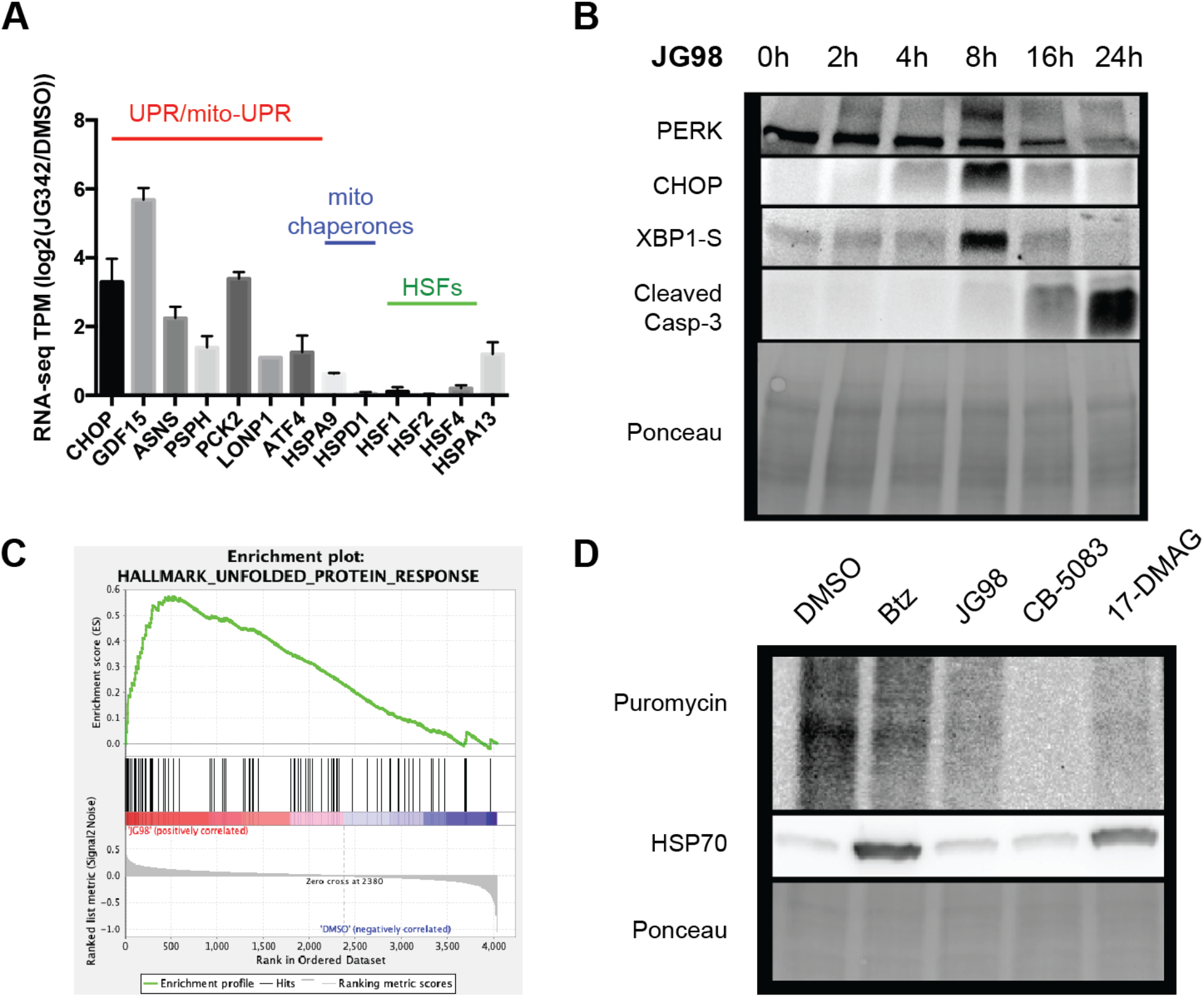
Further investigation of the UPR and translational slowdown. **A.** RNA-seq data for Log_2_-fold changes for JG342 vs. DMSO treated MM.1S cells from 26-hour timepoint (as in **Fig 5E**). **B.** Unfolded Protein Response (UPR) induction in MM.1S cells treated with 3 μM JG98. **C. “** Unfolded Protein Response” has the highest net enrichment in GSEA analysis for JG98 treatment combined proteomic data from three cell lines (Normalized enrichment score (NES) = 2.4, FDR q-value = 0.043, data from experiment outlined in Fig. 4A). **D.** Translational slowdown via puromycin incorporation and total HSP70 levels in KMS34 cells treated with 2μM JG98, 7.5nM Bortezomib, 1μM CB-5083, 500nM 17-DMAG, or DMSO for 22 hours and 1 μM Puromycin for 1 hour. Consistent with proteomic data, only Btz and 17-DMAG show an increase in total HSP70, while all drugs lead to translational slowdown due to the Integrated Stress Response.

**Supplementary Figure 9.**
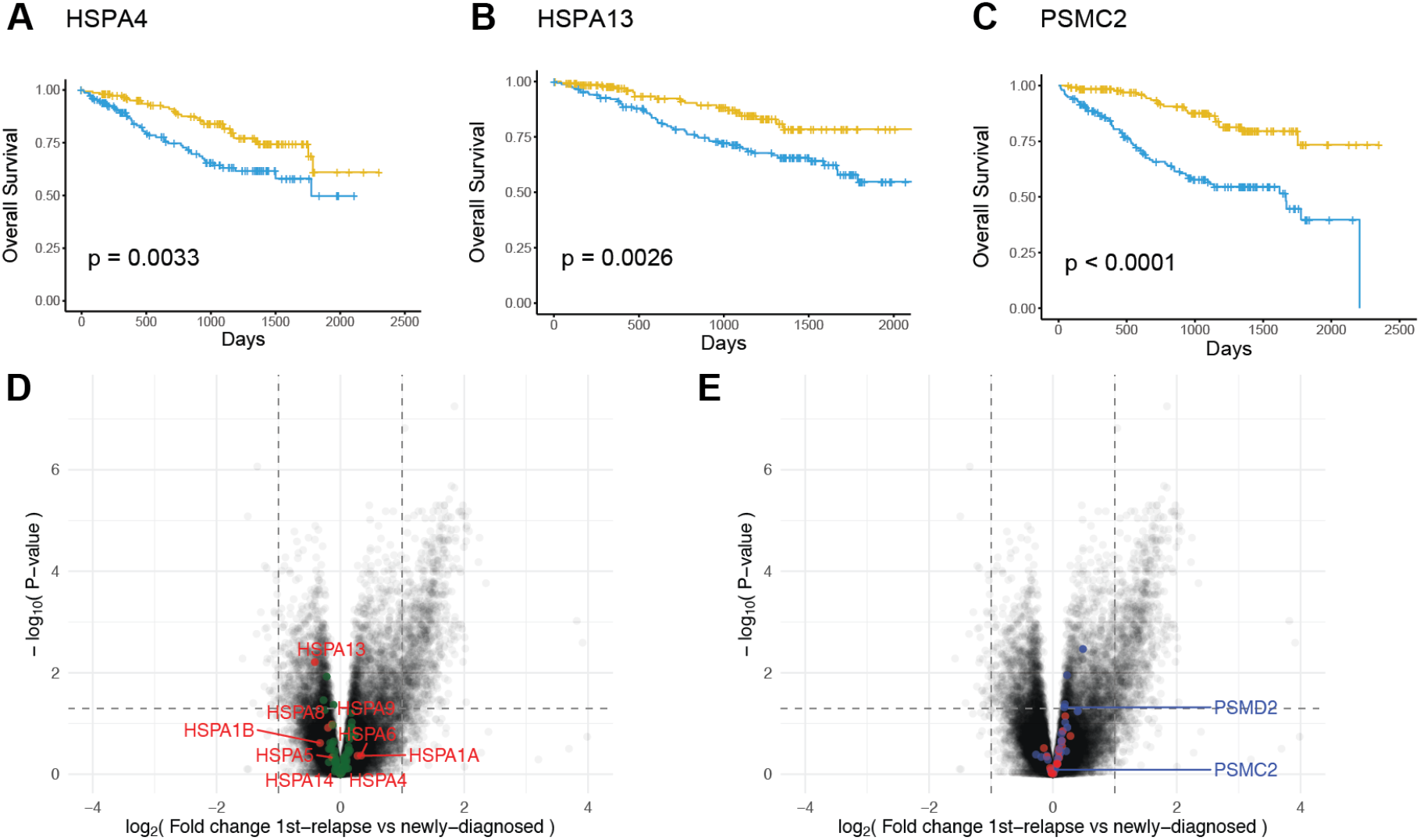
Further survival correlations in MM patients of proteostasis genes and disease evolution at first relapse. **A-C.** Kaplan-Meier curves showing overall survival from CoMMpass trial for patients stratified by *HSPA4*, *HSPA13*, and *PSMC2* RNA levels (patients in top 20% of gene expression in blue, bottom 20% in gold, as in **Fig. 7**). **D-E.** Volcano plot illustrating RNA-seq expression fold changes (in counts) for 50 CoMMpass patients with tumor sample at 1^st^-relapse vs. diagnosis. HSP70 isoforms labeled in red and mitoribosome subunits labeled in green (**D**) and 20S core proteasome subunits labeled in red and 19S cap subunits in blue (**E**).

**Supplementary Table 1.**
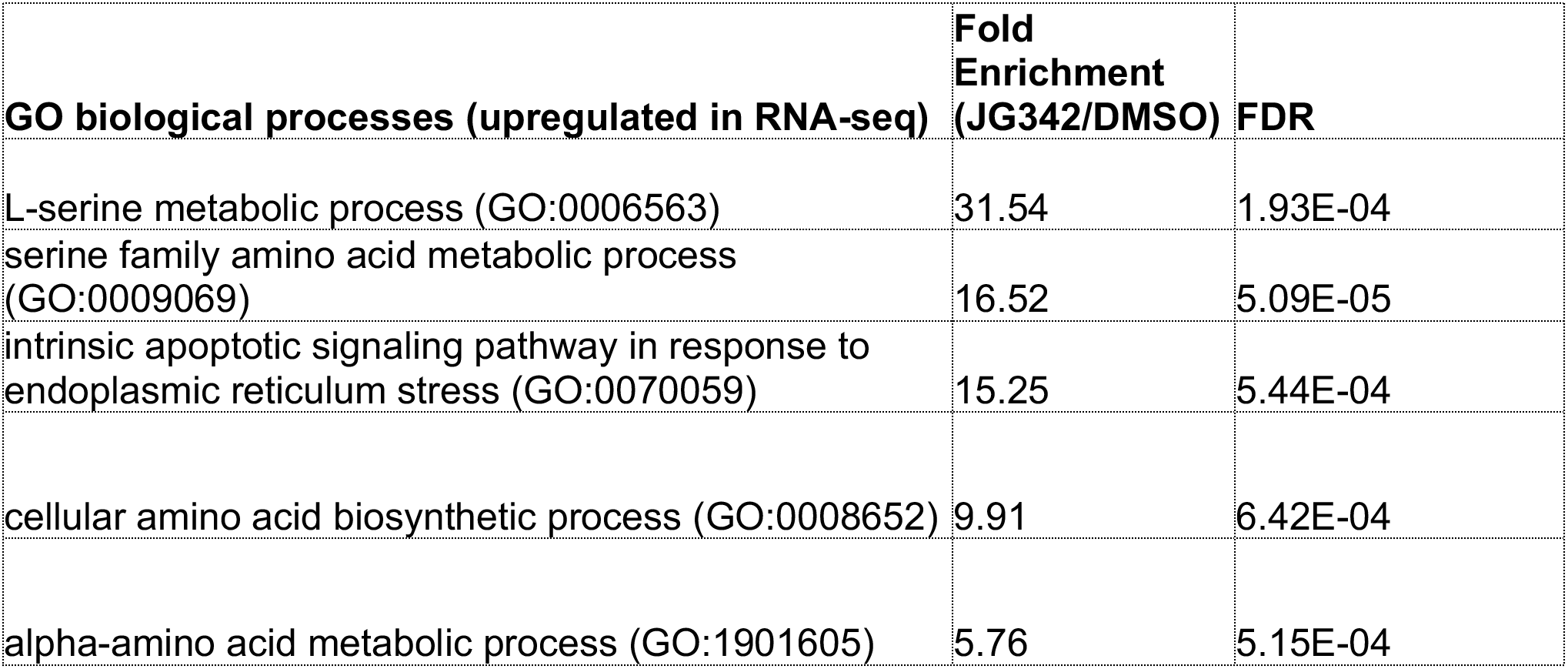
Upregulated Gene Ontology biological processes after JG342 treatment in MM.1S cells from RNA-seq data (FDR <1e-3)

| **GO biological processes (upregulated in RNA-seq)** | **Fold Enrichment (JG342/DMSO)** | **FDR** |
| --- | --- | --- |
| L-serine metabolic process (GO:0006563) | 31.54 | 1.93E-04 |
| serine family amino acid metabolic process (GO:0009069) | 16.52 | 5.09E-05 |
| intrinsic apoptotic signaling pathway in response to endoplasmic reticulum stress (GO:0070059) | 15.25 | 5.44E-04 |
| cellular amino acid biosynthetic process (GO:0008652) | 9.91 | 6.42E-04 |
| alpha-amino acid metabolic process (GO:1901605) | 5.76 | 5.15E-04 |

**Supplementary Dataset 1 (attached as Excel file)**: Curated list of genes included in proteostasis mapping.

**Supplementary Dataset 2 (attached as Excel file):** Proteomic data results from TMT shotgun proteomics experiments.

**Supplementary Dataset 3 (attached as Excel file):** Proteomic data results from TMT-pSILAC proteomics experiments.

## REFERENCES

1. Manasanch, E.E. & Orlowski, R.Z. Proteasome inhibitors in cancer therapy. Nat Rev Clin Oncol 14, 417 (2017).

2. Obeng, E.A. et al. Proteasome inhibitors induce a terminal unfolded protein response in multiple myeloma cells. Blood 107, 4907–4916 (2006).

3. Vincenz, L., Jager, R., O’Dwyer, M. & Samali, A. Endoplasmic reticulum stress and the unfolded protein response: targeting the Achilles heel of multiple myeloma. Mol Cancer Therap 12, 831–843 (2013).

4. Kambhampati, S. & Wiita, A.P. Lessons learned from proteasome inhibitors, the paradigm for targeting protein homeostasis in cancer. Adv Exp Med Biol 1243, 147–162 (2020).

5. Barrio, S. et al. Spectrum and functional validation of PSMB5 mutations in multiple myeloma. Leukemia 33, 447–456 (2019).

6. Farrell, M.L. & Reagan, M.R. Soluble and Cell-Cell-Mediated Drivers of Proteasome Inhibitor Resistance in Multiple Myeloma. Front Endocrinol 9, 218 (2018).

7. Nikesitch, N. & Ling, S.C. Molecular mechanisms in multiple myeloma drug resistance. J Clin Pathol 69, 97–101 (2016).

8. Leung-Hagesteijn, C. et al. Xbp1s-negative tumor B cells and pre-plasmablasts mediate therapeutic proteasome inhibitor resistance in multiple myeloma. Cancer Cell 24, 289–304 (2013).

9. Soriano, G.P. et al. Proteasome inhibitor-adapted myeloma cells are largely independent from proteasome activity and show complex proteomic changes, in particular in redox and energy metabolism. Leukemia (2016).

10. Acosta-Alvear, D. et al. Paradoxical resistance of multiple myeloma to proteasome inhibitors by decreased levels of 19S proteasomal subunits. eLife 4, e08153 (2015).

11. Tsvetkov, P. et al. Compromising the 19S proteasome complex protects cells from reduced flux through the proteasome. eLife 4(2015).

12. Wiita, A.P. et al. Global cellular response to chemotherapy-induced apoptosis. eLife 2, e01236 (2013).

13. Mitsiades, N. et al. Molecular sequelae of proteasome inhibition in human multiple myeloma cells. Proc Natl Acad Sci USA 99, 14374–14379 (2002).

14. Li, X. et al. Validation of the Hsp70-Bag3 protein-protein interaction as a potential therapeutic target in cancer. Mol Cancer Therap 14, 642–648 (2015).

15. Li, X. et al. Analogs of the Allosteric Heat Shock Protein 70 (Hsp70) Inhibitor, MKT-077, as Anti-Cancer Agents. ACS Med Chem Lett 4(2013).

16. Shao, H. et al. Exploration of Benzothiazole Rhodacyanines as Allosteric Inhibitors of Protein-Protein Interactions with Heat Shock Protein 70 (Hsp70). J Med Chem 61, 6163–6177 (2018).

17. Tsherniak, A. et al. Defining a Cancer Dependency Map. Cell 170, 564–576 e516 (2017).

18. Eugenio, A.I.P. et al. Proteasome and heat shock protein 70 (HSP70) inhibitors as therapeutic alternative in multiple myeloma. Oncotarget 8, 114698–114709 (2017).

19. Gabai, V.L. et al. Anticancer Effects of Targeting Hsp70 in Tumor Stromal Cells. Cancer Res 76, 5926–5932 (2016).

20. Moses, M.A. et al. Targeting the Hsp40/Hsp70 Chaperone Axis as a Novel Strategy to Treat Castration-Resistant Prostate Cancer. Cancer Res 78, 4022–4035 (2018).

21. Srinivasan, S.R. et al. Heat Shock Protein 70 (Hsp70) Suppresses RIP1-Dependent Apoptotic and Necroptotic Cascades. Mol Cancer Res 16, 58–68 (2018).

22. Besse, A. et al. Carfilzomib resistance due to ABCB1/MDR1 overexpression is overcome by nelfinavir and lopinavir in multiple myeloma. Leukemia 32, 391–401 (2018).

23. Chitta, K. et al. Targeted inhibition of the deubiquitinating enzymes, USP14 and UCHL5, induces proteotoxic stress and apoptosis in Waldenstrom macroglobulinaemia tumour cells. Br J Haematol 169, 377–390 (2015).

24. Das, D.S. et al. Blockade of Deubiquitylating Enzyme USP1 Inhibits DNA Repair and Triggers Apoptosis in Multiple Myeloma Cells. Clin Cancer Res 23, 4280 (2017).

25. Le Moigne, R. et al. The p97 inhibitor CB-5083 is a unique disrupter of protein homeostasis in models of Multiple Myeloma. Mol Cancer Therap 16, 2375 (2017).

26. Tian, Z. et al. A novel small molecule inhibitor of deubiquitylating enzyme USP14 and UCHL5 induces apoptosis in multiple myeloma and overcomes bortezomib resistance. Blood 123, 706–716 (2014).

27. Mitra, A.K. et al. A gene expression signature distinguishes innate response and resistance to proteasome inhibitors in multiple myeloma. Blood Cancer J 7, e581 (2017).

28. Egorin, M.J. et al. Pharmacokinetics, tissue distribution, and metabolism of 17-(dimethylaminoethylamino)-17-demethoxygeldanamycin (NSC 707545) in CD2F1 mice and Fischer 344 rats. Cancer Chemotherap Pharamcol 49, 7–19 (2002).

29. Anderson, D.J. et al. Targeting the AAA ATPase p97 as an Approach to Treat Cancer through Disruption of Protein Homeostasis. Cancer Cell 28, 653–665 (2015).

30. Yadav, B., Wennerberg, K., Aittokallio, T. & Tang, J. Searching for Drug Synergy in Complex Dose-Response Landscapes Using an Interaction Potency Model. Comp Struct Biotechnol J 13, 504–513 (2015).

31. Modica-Napolitano, J.S. & Aprille, J.R. Delocalized lipophilic cations selectively target the mitochondria of carcinoma cells. Adv Drug Deliv Rev 49, 63–70 (2001).

32. Wadhwa, R. et al. Selective toxicity of MKT-077 to cancer cells is mediated by its binding to the hsp70 family protein mot-2 and reactivation of p53 function. Cancer Res 60, 6818–6821 (2000).

33. Shao, H. & Gestwicki, J.E. Neutral analogs of the heat shock protein 70 (Hsp70) inhibitor, JG-98. Bioorg Med Chem Lett 30, 126954 (2020).

34. Wu, P.K. et al. Mortalin (HSPA9) facilitates BRAF-mutant tumor cell survival by suppressing ANT3-mediated mitochondrial membrane permeability. Sci Signaling 13(2020).

35. Ramkumar, P. et al. CRISPR-based screens uncover determinants of immunotherapy response and potential combination therapy strategies. BioRxiv doi: 10.1101/833707 (2019).

36. Besse, L. et al. A metabolic switch in proteasome inhibitor-resistant multiple myeloma ensures higher mitochondrial metabolism, protein folding and sphingomyelin synthesis. Haematologica 104, e415–e419 (2019).

37. Tsvetkov, P. et al. Mitochondrial metabolism promotes adaptation to proteotoxic stress. Nat Chem Biol 15, 681–689 (2019).

38. Rosenzweig, R., Nillegoda, N.B., Mayer, M.P. & Bukau, B. The Hsp70 chaperone network. Nat Rev Mol Cell Biol 20, 665–680 (2019).

39. Savitski, M.M. et al. Multiplexed Proteome Dynamics Profiling Reveals Mechanisms Controlling Protein Homeostasis. Cell 173, 260–274 e225 (2018).

40. Welle, K.A. et al. Time-resolved Analysis of Proteome Dynamics by Tandem Mass Tags and Stable Isotope Labeling in Cell Culture (TMT-SILAC) Hyperplexing. Mol Cell Proteom 15, 3551–3563 (2016).

41. Hoedt, E., Zhang, G. & Neubert, T.A. Stable Isotope Labeling by Amino Acids in Cell Culture (SILAC) for Quantitative Proteomics. Adv Exp Med Biol 1140, 531–539 (2019).

42. Pakos-Zebrucka, K. et al. The integrated stress response. EMBO Rep 17, 1374–1395 (2016).

43. Sidrauski, C., McGeachy, A.M., Ingolia, N.T. & Walter, P. The small molecule ISRIB reverses the effects of eIF2alpha phosphorylation on translation and stress granule assembly. eLife 4(2015).

44. Candas, D. & Li, J.J. MnSOD in oxidative stress response-potential regulation via mitochondrial protein influx. Antiox Redox Signaling 20, 1599–1617 (2014).

45. Valera-Alberni, M. & Canto, C. Mitochondrial stress management: a dynamic journey. Cell Stress 2, 253–274 (2018).

46. Sabnis, A.J. et al. Combined chemical-genetic approach identifies cytosolic HSP70 dependence in rhabdomyosarcoma. Proc Natl Acad Sci USA 113, 9015–9020 (2016).

47. Walter, P. & Ron, D. The unfolded protein response: from stress pathway to homeostatic regulation. Science 334, 1081–1086 (2011).

48. Mulligan, G. et al. Gene expression profiling and correlation with outcome in clinical trials of the proteasome inhibitor bortezomib. Blood 109, 3177–3188 (2007).

49. Shah, S.P., Lonial, S. & Boise, L.H. When Cancer Fights Back: Multiple Myeloma, Proteasome Inhibition, and the Heat-Shock Response. Mol Cancer Res 13, 1163–1173 (2015).

50. Shah, S.P. et al. Bortezomib-induced heat shock response protects multiple myeloma cells and is activated by heat shock factor 1 serine 326 phosphorylation. Oncotarget 7, 59727–59741 (2016).

51. Fontaine, S.N. et al. Isoform-selective Genetic Inhibition of Constitutive Cytosolic Hsp70 Activity Promotes Client Tau Degradation Using an Altered Co-chaperone Complement. J Biol Chem 290, 13115–13127 (2015).

52. Taguwa, S. et al. Defining Hsp70 Subnetworks in Dengue Virus Replication Reveals Key Vulnerability in Flavivirus Infection. Cell 163, 1108–1123 (2015).

53. Love, M.I., Huber, W. & Anders, S. Moderated estimation of fold change and dispersion for RNA-seq data with DESeq2. Genome Biol 15, 550 (2014).

54. Barretina, J. et al. The Cancer Cell Line Encyclopedia enables predictive modelling of anticancer drug sensitivity. Nature 483, 603–607 (2012).

55. Mitra, A.K. et al. Single-cell analysis of targeted transcriptome predicts drug sensitivity of single cells within human myeloma tumors. Leukemia 30, 1094–1102 (2016).

56. Perelman, A. et al. JC-1: alternative excitation wavelengths facilitate mitochondrial membrane potential cytometry. Cell Death Dis 3, e430 (2012).

57. Miyata, Y. et al. Synthesis and initial evaluation of YM-08, a blood-brain barrier permeable derivative of the heat shock protein 70 (Hsp70) inhibitor MKT-077, which reduces tau levels. ACS Chem Neurosci 4, 930–939 (2013).

58. Tian, R. et al. CRISPR Interference-Based Platform for Multimodal Genetic Screens in Human iPSC-Derived Neurons. Neuron 104, 239 (2019).

59. Kim, D., Langmead, B. & Salzberg, S.L. HISAT: a fast spliced aligner with low memory requirements. Nat Methods 12, 357–360 (2015).

60. Langmead, B. & Salzberg, S.L. Fast gapped-read alignment with Bowtie 2. Nat Methods 9, 357–359 (2012).

61. Li, B. & Dewey, C.N. RSEM: accurate transcript quantification from RNA-Seq data with or without a reference genome. BMC Bioinformatics 12, 323 (2011).

62. Huang, H.H. et al. Proteasome inhibitor-induced modulation reveals the spliceosome as a specific therapeutic vulnerability in multiple myeloma. Nat Commun in press(2020).

63. Zecha, J. et al. TMT Labeling for the Masses: A Robust and Cost-efficient, In-solution Labeling Approach. Mol Cell Proteom 18, 1468–1478 (2019).

64. Tyanova, S., Temu, T. & Cox, J. The MaxQuant computational platform for mass spectrometry-based shotgun proteomics. Nat Protoc 11, 2301–2319 (2016).

65. Tyanova, S. et al. The Perseus computational platform for comprehensive analysis of (prote)omics data. Nat Methods 13, 731–740 (2016).

66. Perez-Riverol, Y. et al. The PRIDE database and related tools and resources in 2019: improving support for quantification data. Nuc Acid Res 47, D442–D450 (2019).

67. Subramanian, A. et al. Gene set enrichment analysis: a knowledge-based approach for interpreting genome-wide expression profiles. Proc Natl Acad Sci USA 102, 15545–15550 (2005).

